# Quantifying the spatiotemporal dynamics of IRES versus Cap translation with single-molecule resolution in living cells

**DOI:** 10.1101/2020.01.09.900829

**Authors:** Amanda Koch, Luis Aguilera, Tatsuya Morisaki, Brian Munsky, Timothy J. Stasevich

## Abstract

Viruses use IRES sequences within their RNA to hijack translation machinery and thereby rapidly replicate in host cells. While this process has been extensively studied in bulk assays, the dynamics of hijacking at the single-molecule level remain unexplored in living cells. To achieve this, we developed a bicistronic biosensor encoding complementary repeat epitopes in two ORFs, one translated in a Cap-dependent manner and the other translated in an IRES-mediated manner. Using a pair of complementary probes that bind the epitopes co-translationally, our biosensor lights up in different colors depending on which ORF is being translated. In combination with single-molecule tracking and computational modeling, we measured the relative kinetics of Cap versus IRES translation and show: (1) Two non-overlapping ORFs can be simultaneously translated within a single mRNA; (2) EMCV IRES-mediated translation sites recruit ribosomes less efficiently than Cap-dependent translation sites but are otherwise nearly indistinguishable, having similar mobilities, sizes, spatial distributions, and ribosomal initiation and elongation rates; (3) Both Cap-dependent and IRES-mediated ribosomes tend to stretch out translation sites; (4) Although the IRES recruits two to three times fewer ribosomes than the Cap in normal conditions, the balance shifts dramatically in favor of the IRES during oxidative and ER stresses that mimic viral infection; and (5) Translation of the IRES is enhanced by translation of the Cap, demonstrating upstream translation can positively impact the downstream translation of a non-overlapping ORF. With the ability to simultaneously quantify two distinct translation mechanisms in physiologically relevant live-cell environments, we anticipate bicistronic biosensors like the one we developed here will become powerful new tools to dissect both canonical and non-canonical translation dynamics with single-molecule precision.

**Graphical Abstract:** 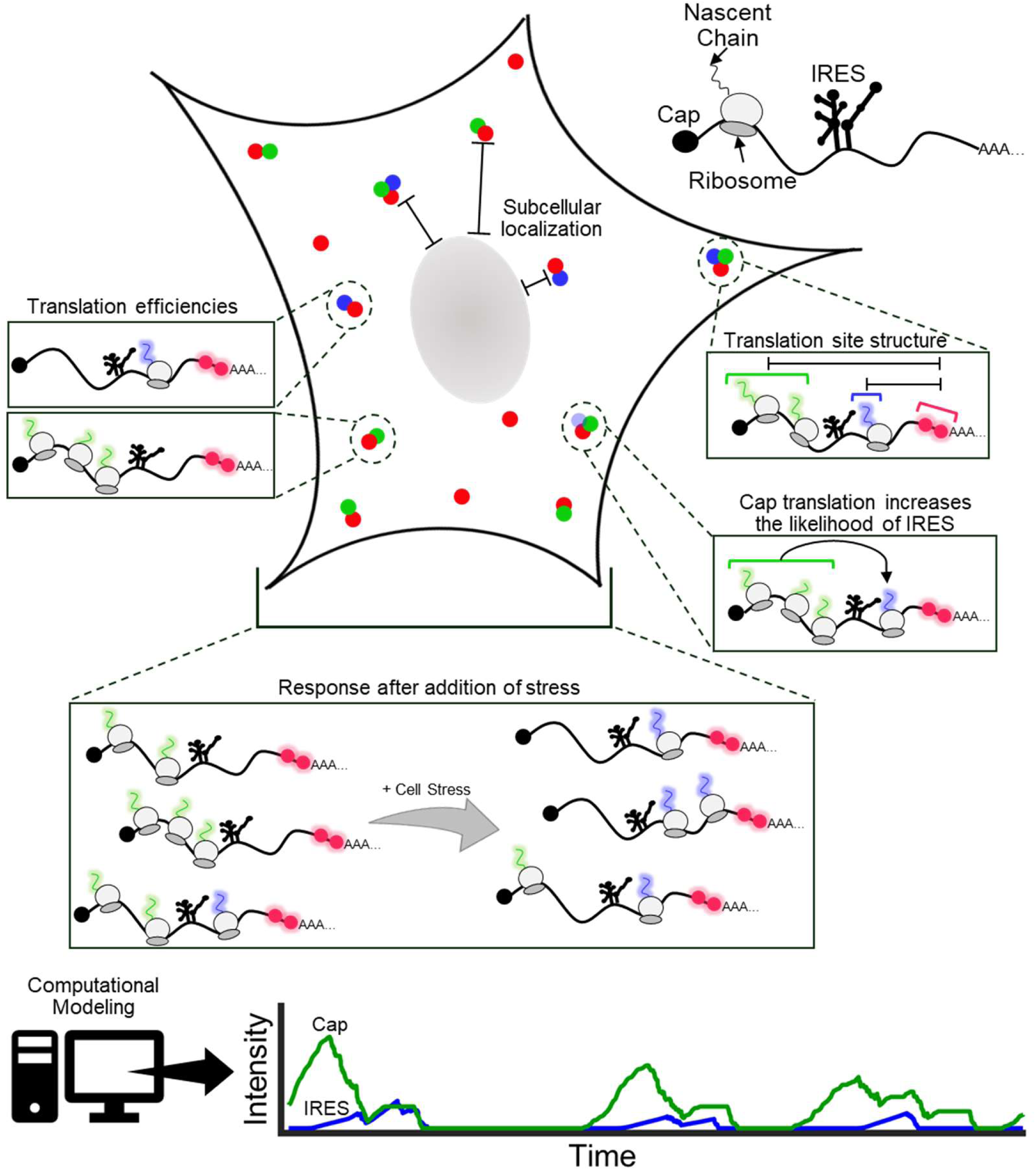

## INTRODUCTION

Viruses are efficient parasites in part because they have evolved non-canonical mechanisms to hijack host translation machinery to their benefit.^1, 2^ Internal ribosome entry sites (IRES) within the 5’ untranslated regions of viral RNA, for example, act as lures that attract host ribosomes and activate viral translation without the need for the canonical 5’ Cap most host transcripts harbor.^2–7^

To achieve this, IRES sequences have evolved into several distinct classes, each containing unique 2D and 3D structural RNA motifs^4, 8^ that attract different subsets of host translation factors to varying degrees.^2, 4–7^ The entire process is furthermore sped up by the active induction of cellular stress pathways upon viral infection that generally repress canonical Cap-dependent translation.^9–14^ The end result is a large pool of host initiation factors and ribosomal subunits that are free to bind and initiate at viral IRES sequences at the peril of the host cell.^15–18^

Previous experimental techniques used to measure the efficiency of IRES elements involve the use of a bicistronic transcript encoding an IRES between two reporter proteins.^19, 20^ The construct is transiently transfected in cells, and the relative expression of the two reporters is later quantified, often in bulk via fluorescent activated cell sorting (FACS) ^21^ or a dual-luciferase assay.^8^ In either case, the IRES activity is determined by the ratio of expressed reporter downstream of the IRES versus expressed reporter upstream of the IRES.^22^ These types of assays have been beneficial for deducing the relative IRES activity in cells hours or days after transfection.^23^ However, they lack the spatiotemporal resolution needed to visualize the sites of IRES translation and quantify translation initiation and elongation kinetics at those sites in real-time. This has made it difficult to assess the heterogeneity of IRES-mediated translation within cells or to see if there is variation between RNA or within specific subcellular environments. Methods to study IRES-mediated translation with higher spatiotemporal resolution are therefore needed to better understand how viruses exploit IRES sequences during viral biogenesis.

Here, we develop and apply a real-time biosensor to quantify IRES-mediated translation dynamics with single-molecule resolution in living cells. To achieve this, we engineered complementary repeat epitopes into a bicistronic reporter, such that Cap-versus IRES-translation could be monitored in two colors simultaneously from a single RNA using Nascent Chain Tracking (NCT).^24^ The resulting biosensor allows us to fairly and accurately measure the dynamics of Cap-versus IRES- translation with spatial resolution on the tens of nanometers scale and temporal resolution on the sub-seconds timescale. Application to the IRES from the Encephalomyocarditis Virus (EMCV) uncovers the spatial organization and dynamics of single IRES-mediated translation sites compared to those driven by the Cap under both normal and stressful physiological conditions. Given the ubiquity of viral translation, we anticipate our biosensor will find broad application, not just to better understand viral replication, but also in screens to help identify and develop novel therapeutics that selectively perturb IRES-mediated translation.

## RESULTS

### A multicolor biosensor to compare Cap and IRES translation at the single-molecule level in living cells

To analyze the translation initiation mechanisms of a viral IRES compared to canonical Cap, we constructed a bicistronic NCT biosensor that is bound and lit up by different fluorescent probes depending on the manner of translation initiation (Figure 1A-B). Encoded under Cap-dependent translation is a lysine demethylase KDM5B, N-terminally fused to a spaghetti monster tag (SM) encoding 10×FLAG epitopes. The FLAG SM tag is bound by fluorescently conjugated fragments of anti-FLAG antibodies (Fab), allowing us to begin visualization of Cap translation soon after the first FLAG epitope emerges from the ribosome.^24, 25^ Encoded under IRES-mediated translation is a Kinesin-like protein Kif18b, N-terminally tagged with 24×SunTag epitopes.^26^ The SunTag epitopes are bound by single chain variable fragments (scFv) fused to GFP. As with FLAG SM, visualization of IRES-mediated translation occurs soon after the first SunTag epitope emerges from the ribosome.^24, 26^ In addition, the biosensor contains 24×MS2 stem loops in the 3’ untranslated region (UTR) which, upon transcription, are bound by Halo-tagged MS2 coat proteins (MCP) (Figure 1A-B).^25^ The construct allows for the simultaneous imaging of Cap-dependent and IRES-mediated translation from single transcripts.

**Figure 1:**
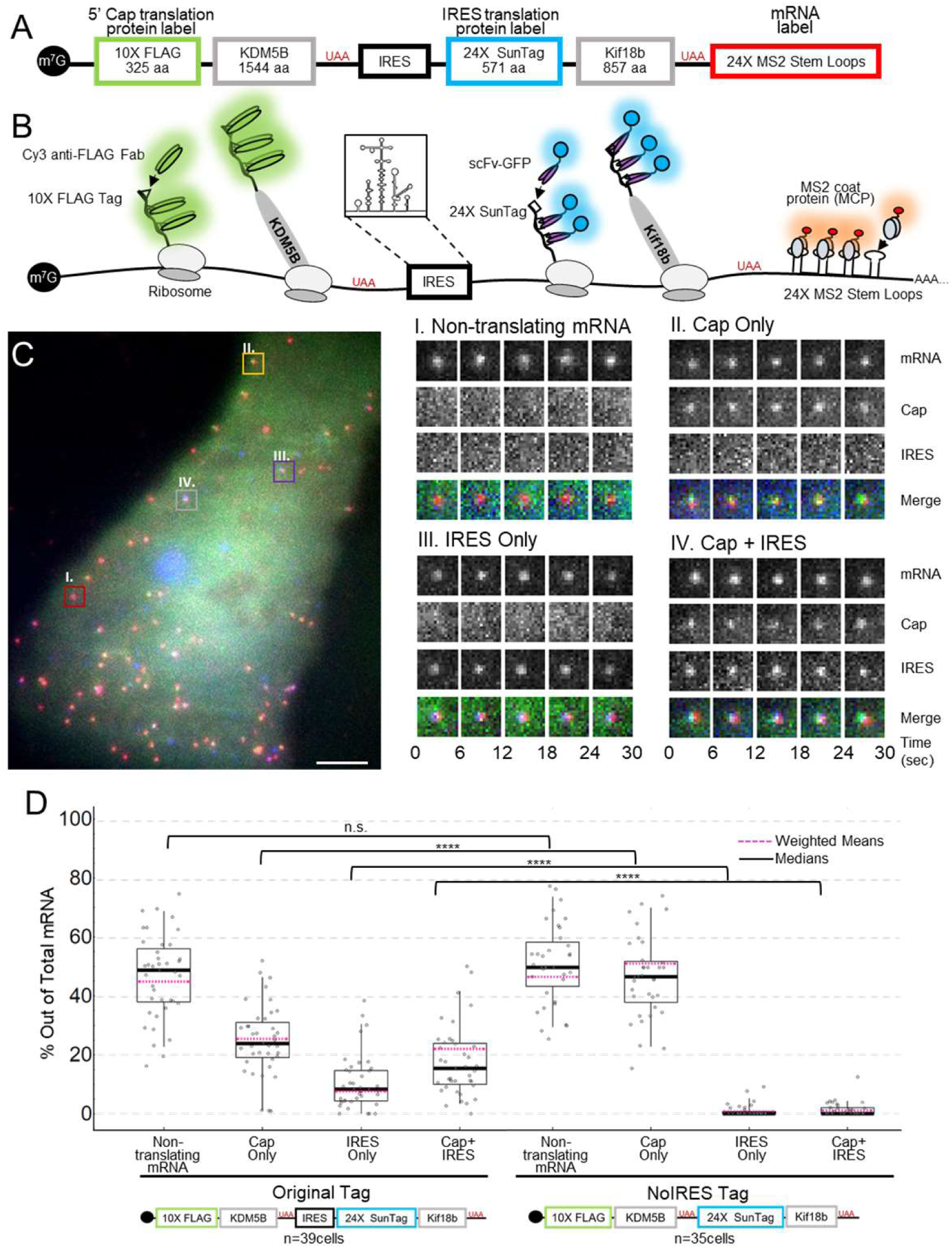
A multicolor biosensor to compare Cap and IRES translation at the single-molecule level in living cells. (A) Plasmid encodes KDM5B with an N-terminal 10xFLAG SM tag under Cap-dependent translation initiation and Kif18b with an N-terminal 24xSuntag under IRES-mediated translation initiation. Between KDM5B and SunTag is encoded the full EMCV IRES element. Encoded in the 3’ UTR are 24xMS2 stem loops. (B) Schematic of the system. RNA (red) is marked by MCP-Halo labeled with JF646 that binds to repeated MS2 stem loops in the 3’ UTR. Cap-dependent protein reporter (green) is labeled by anti-FLAG Fab conjugated to Cy3 that binds the 10× FLAG peptide epitopes in the N-terminus. IRES-mediated protein reporter (blue) is labeled by a GCN4 scFv fused to a GFP that binds the 24× SunTag peptide epitopes. (C) Representative cell imaged 6 hours after plasmid and probe loading. Different colored boxes within the cell illustrate different types of translation spots seen within a single cell. Example co-moving spots to right of cell. I – non-translating mRNA (red). II – single mRNA translating Cap Only (yellow). III – single mRNA translating IRES Only (purple). IV – single mRNA translating in a Cap and IRES manner (gray). (D) Quantification of species percentages out of total mRNA for both the Original Tag and the noIRES control. Each point represents the percent of that species in a cell. The p-values are based on a two-tailed Mann-Whitney test: *p<0.05, **p<0.01, ***p<0.001, ****p<0.0001. The thick black line indicate the median and the dashed red line represents the weighted (by mRNA/cell) mean, the box indicate the 25%-75% range, and the whiskers indicate the 5%-95% range.

As a first application, we inserted the IRES element from EMCV into our biosensor. EMCV is a small single-stranded RNA virus that has been shown to cause many diseases in mammals.^27^ The IRES sequence that this virus uses to replicate is 553 nucleotides in length and contains a methionine start codon that defines the start of the preferred open reading frame (ORF) (Figure 1B).^28, 29^ Previous assays have shown this IRES can recruit ribosomes without the need of a 5’ Cap or many characterized canonical translation initiation factors.^28, 29^ Because of these properties, the EMCV IRES is widely used in research settings to coexpress distinct gene products from a single transcript.^28, 29^ However, little is known about the intracellular localization, mobility, and general translation efficiency of the EMCV IRES in living cells at the single-molecule level.

To begin to visualize these aspects of EMCV IRES translation, we transiently transfected our biosensor into living cells, and 3-6 hours later we bead loaded anti-FLAG Cy3-Fab, purified anti-SunTag GFP-scFv, and purified HaloTag-MCP. With this combination of probes, translation sites could be identified by protein labeled by Fab or scFv co-moving with mRNA labeled by MCP. In addition to non-translating mRNA (Figure 1C panel I), we were able to identify translation sites labeled by just Fab (Figure 1C panel II), just scFv (Figure 1C panel III), and both Fab and scFv (Figure 1C panel IV), indicating Cap-only translation, IRES-only translation, and Cap+IRES translation, respectively.

We performed two control studies to confirm that these spots were active translation sites. First, to rule out the possibility of fluorescence bleed-through from one of the protein channels to the RNA channel, we repeated experiments without labeling RNA. Consistent with no bleed-through, we did not observe detectable fluorescence in the RNA channel, regardless of the intensity of translation sites (Figure S1A-B). All other forms of bleed-through were ruled out by direct observations of distinct populations of non-translating mRNA, IRES-only translation sites, and Cap-only translation sites. Second, to confirm that the translation sites were active, we treated cells with puromycin, an elongation inhibitor that releases nascent chains from ribosomes.^30^ Consistent with this action, upon addition of 50 ug/mL puromycin, we observed the rapid disappearance of all Fab/scFv translation signals within translation sites (Figure S1C).

To get a better sense of the heterogeneity of translation sites we could observe with our system, we took 2.5 minute movies (25 frames × 13 z-planes per volume × 3 colors = 975 images per movie) of 39 cells expressing our biosensor. Movies were acquired over eight independent imaging days using oblique HILO illumination to maximize signal-to-noise.^31^ In total, we measured translation dynamics from 3748 mRNA, of which 1784 were being translated. Within cells, a median of 24% of mRNA were translated in a Cap-only manner, 8% in an IRES-only manner, and 15% in both a Cap- and IRES-manner simultaneously (Cap+IRES) (Figure 1D, left). As a control, we removed the IRES element from our biosensor. In this case, we observed virtually no sites being translated in an IRES-mediated manner (Figure 1D, right), ruling out the possibility of ribosomal run-through from Cap to IRES, as well as the presence of an unknown splicing donor site or a cryptic initiation site. All else equal, these data demonstrate the IRES element alone can capture and initiate host ribosomes, but not as efficiently as the Cap. Given the relatively large fraction of transcripts we observed being translated in both a Cap- and IRES- manner, these data also demonstrate that a single bicistronic transcript can indeed have two open reading frames translated simultaneously.

### IRES and Cap translation sites localize and move similarly within cells

Since we observed large pools of both Cap-dependent and IRES-mediated translation sites in our cells, we wondered if any biophysical properties of the sites correlated with their mode of translation. We first checked to see if translation sites have a tendency to cluster. For this, we measured the distance from each mRNA to the nearest neighbor translation site. Compared to non-translating mRNA, translating mRNA tended to be significantly closer to other translation sites, but this was true regardless of whether or not translation occurred in a Cap-dependent or IRES-mediated manner (Figure 2A). Next, we measured the shortest distance between each mRNA and the edge of the nucleus (Figure 2B). According to this metric, Cap-only translation sites were statistically indistinguishable from Cap+IRES translation sites and non-translating mRNA, while IRES-only translation sites had a slight preference for the perinuclear region.

**Figure 2:**
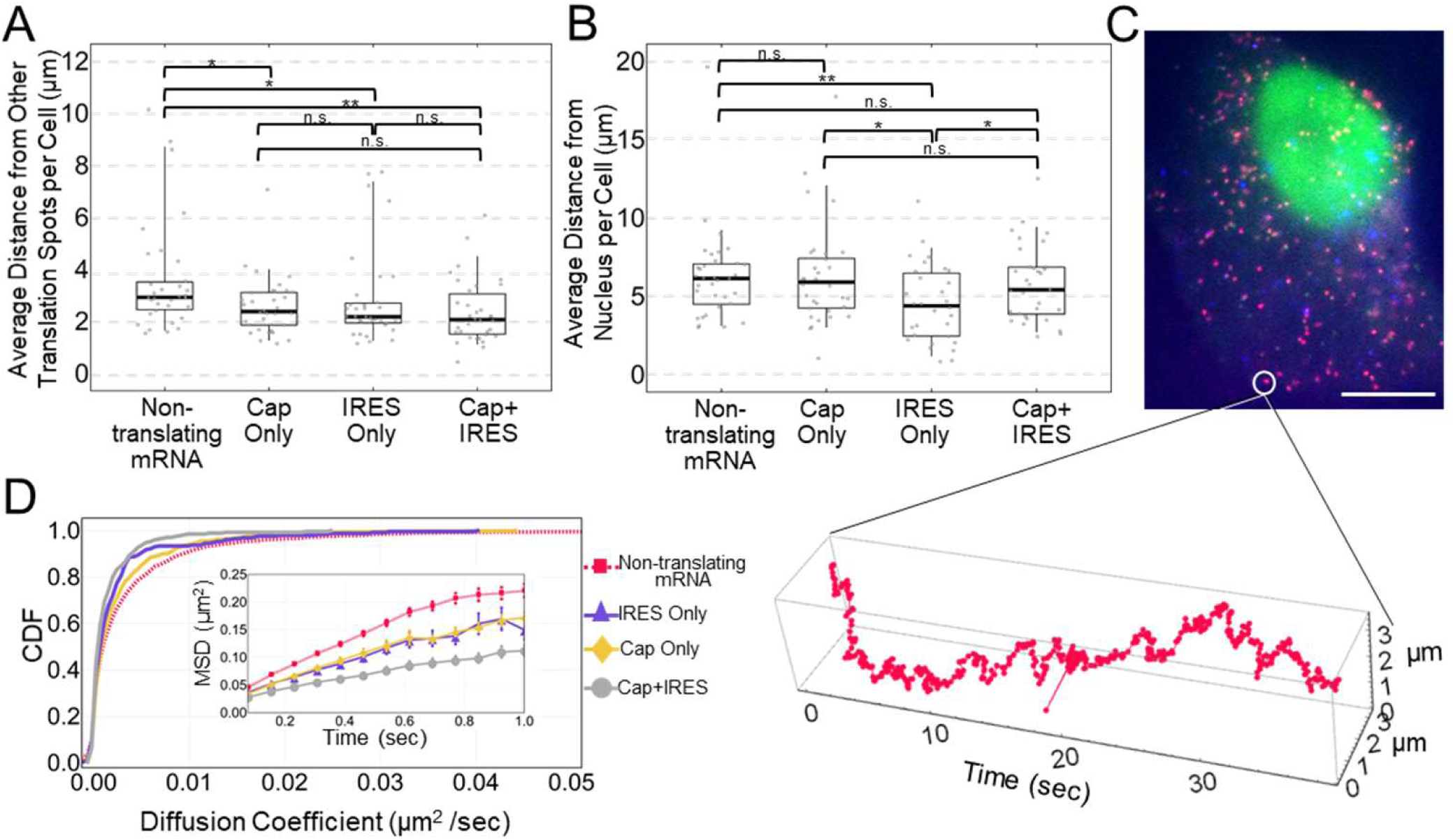
IRES and Cap translation sites localize and move similarly within cells. (A) Quantification of translating and non-translating mRNA distances in micrometers (µm) to nearest-neighbor translation spot within single cells. Each point represents the average distance per cell. (B) Quantification of distance in µm from the nucleus of translating and non-translating mRNA. Each point represents the average distance from the edge of the nucleus per cell. (C) Representative cell imaged with fast imaging conditions. An example mRNA is highlighted with a white circle and a track through time of that mRNA is graphed below. (D) Cumulative distribution function plot of non-translating mRNA (red), IRES Only (purple), Cap Only (yellow), and Cap+IRES (gray) species based on their diffusion coefficients (µm^2^ /sec). Inset shows the Mean Square Displacements (MSD) of the different species over time in seconds. n=3771 total tracked mRNA (translating and non-translating), n=11 cells.

Since IRES-only translation sites were slightly closer to the nucleus, we hypothesized these sites may have a unique mobility compared to other translation sites. Indeed, translation sites localized to the perinuclear ER have recently been shown to be less mobile.^32, 33^ To test our hypothesis, we repeated experiments, but now tracking a single plane with a temporal resolution of 77 msec so that even the fastest moving mRNA could be monitored (Figure 2C). From the 3771 mRNA we tracked in this way, we calculated the cumulative distribution of diffusion coefficients, as well as the average mean squared displacement (Figure 2D). In contrast to our hypothesis, the mobility of IRES-only translation sites was not statistically different than Cap-only translation sites. Rather, both IRES-only and Cap-only translation sites displayed intermediate mobility, being on average faster than Cap+IRES translation sites but slower than non-translating mRNA. This overall trend suggests the mobility of our biosensor is mainly dictated by the degree of translation rather than the type of translation, with more ribosomes generally leading to slower overall mobility.^34^

To further test if IRES-mediated translation sites have unique mobilities, we measured their degree of confinement. Recently, large macromolecular complexes such as mRNPs have been shown to exhibit confined movement due to crowding in the cytoplasm.^35^ Confinement generally becomes more severe as the size of the macromolecular complex increases and can be quantified as a preference for 180 degree jumps over 0 degree jumps.^36^ Interestingly, irrespective of translation status, all mRNA in cells displayed some level of confinement (Figure S2). However, again in contrast to our hypothesis, IRES-only translation sites did not show more confinement than Cap-only translation sites. Instead, the degree of confinement was proportional to diffusivity, with Cap+IRES translation sites the most confined and slowest and non-translating mRNA the least confined and fastest. We therefore conclude that the movement and localization of IRES-mediated translation sites is very similar to Cap-dependent translation sites within living cells. Additional information is thus required to accurately distinguish the two types of translation sites.

### IRES and Cap translation sites stretch out as ribosomes load

One of the unique features of our biosensor is that each ORF is long, so Cap-dependent ribosomes on the 5’ end of the biosensor are on average over 5 kb from IRES-mediated ribosomes on the 3’ end and over 8.5 kb from the MS2 signal marking the 3’ UTR (Figure 1A and 3A). This allowed us to accurately measure differences in the positioning of IRES-mediated versus Cap-dependent ribosomes within single translation sites.

Recently, the Zenklusen^37^ and Parker^38^ laboratories used smFISH to show that translating ribosomes tend to stretch out translation sites, i.e. the more strongly an mRNA is translated, the more stretched out it becomes, in contrast to classic models of stable mRNA looping.^39^ To see if we could recapitulate this result in unfixed living cells, we measured the distance between Cap, IRES, and MS2 signals within single-mRNA translation sites over time (Figure 3A-B). We began by analyzing Cap+IRES translation sites because they have two modes of translation and therefore two ways to quantify mRNA stretching (Figure 3B). If an actively translating mRNA were to be stretched out, we would predict spatial separation between all three signals, with IRES signals being closer to the 3’UTR than Cap signals. Consistent with this, the median distance between Cap-dependent ribosomes and the 3’UTR (marked by MCP) was 146 nm, while the median distance between IRES-mediated ribosomes and the 3’UTR was significantly smaller at just 101 nm (Figure 3C, compare top to bottom, p-value = 6E-6 and Figure S3). Furthermore, in agreement with the data from Zenklusen and Parker, by ranking Cap+IRES translation sites by their total intensity (i.e. total ribosomal content or degree of translation), we found that as the brightness of the Cap+IRES signals increased, their distances from the MS2 signal marking the 3’ UTR also generally increased (Figure 3C).

**Figure 3:**
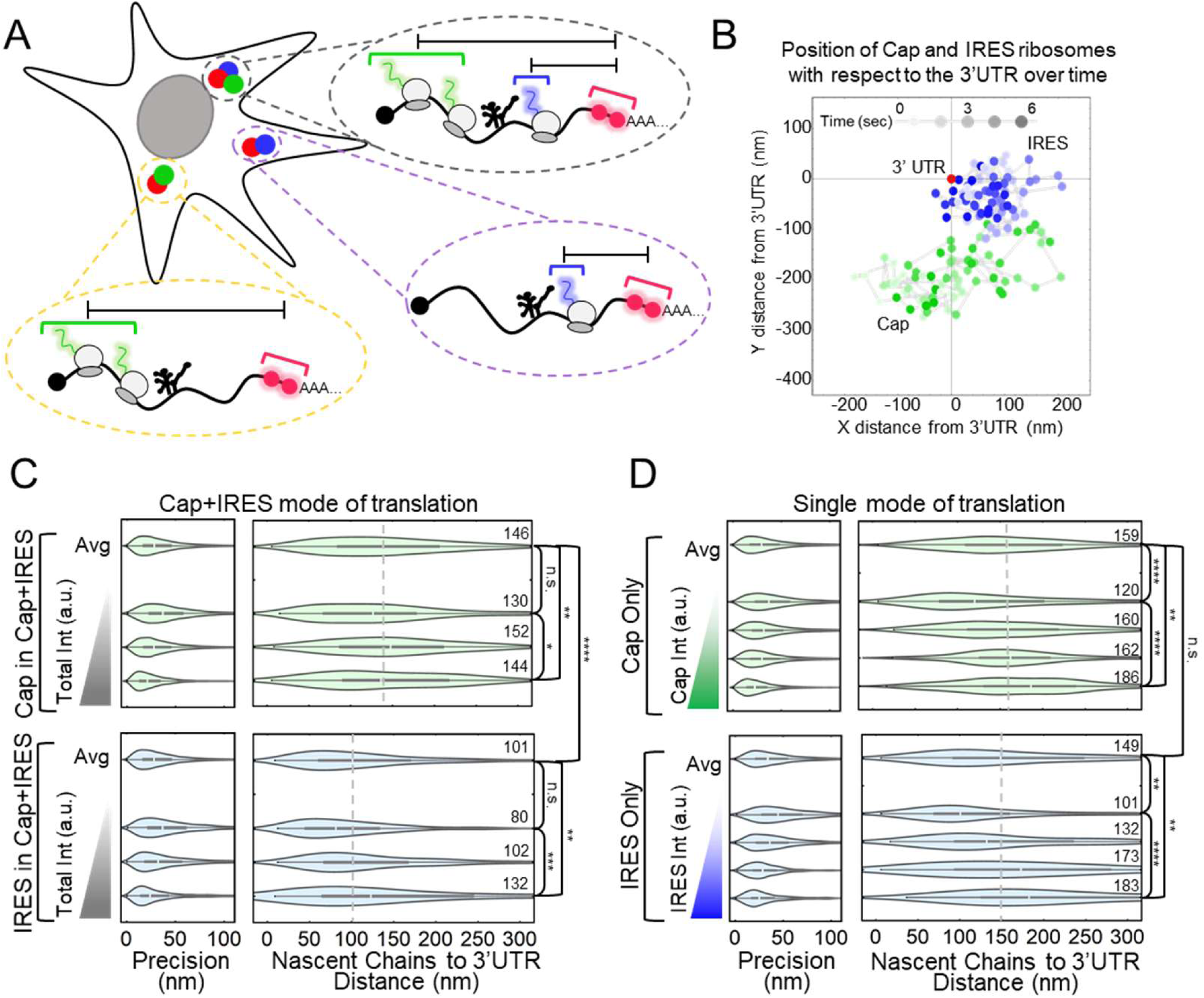
IRES and Cap translation sites stretch out as ribosomes load. (A) Schematic showing how the measurements from the 3’UTR to the Cap and IRES nascent chains were conducted within single cells. (B) Graph showing IRES and Cap nascent chain positions relative to 3’UTR through time of a representative Cap+IRES translation spot. X and Y distances displayed in nanometers (nm) and time (sec) is represented as a gradient in spot color. 75 frames were imaged at a rate of 77 msec per frame. (C) Distributions of the distance between Cap (upper) or IRES (lower) nascent chains and the 3’UTR in Cap+IRES translation sites n=296 translation sites for Cap in Cap+IRES. n=259 translation sites for IRES in Cap+IRES. In each box, the average of all translation sites is shown on top (Avg), and equal-sized subsets sorted by their total nascent chain signal intensity below. Precision is estimated from the distribution of distances between two consecutive timepoints, e.g. any two connected points in B. The median distance is reported to the right of each distribution. (D) Same as (C), but for Cap Only translation sites (upper) n= 793 translation sites and IRES Only translation sites (lower) n=213 translation sites. For the box and whisker plots, the white lines indicate the medians, the boxes indicate the 25%-75% range, and the whiskers indicate the 5%-95% range.

Cap+IRES translation sites have two modes of translation, so it is difficult to know if the stretching within translation sites is primarily due to Cap-dependent ribosomes, IRES-mediated ribosomes, or some combination of the two. To resolve this issue, we next focused on Cap-only and IRES-only translation sites (Figure 3D). Performing the same analysis as before revealed both types of translation sites are significantly stretched out. Specifically, the median distance between Cap-dependent ribosomes and the 3’UTR in Cap-only sites was 159 nm, while the median distance between IRES-mediated ribosomes and the 3’UTR in IRES-only translation sites was slightly smaller at 149 nm (Figure 3D, compare top to bottom, p-value 0.42). Although these single-mode translation sites were a little more stretched out than Cap+IRES translation sites, within single-mode sites the overall trend remained, with the distance between the 3’UTR and the Cap or IRES signals steadily increasing as the degree of translation increased, once again confirming the data from Zenklusen and Parker. We therefore conclude that both IRES-mediated and Cap-dependent translation sites tend to stretch out as ribosomal content increases.

### Elongation does not alter the translation efficiency of the IRES

Despite our inability to detect any clear distinguishing features that predict IRES-mediated translation, the simple observation that it is rare compared to Cap-dependent translation (Figure 1D) suggested either the IRES is less efficient than the Cap at recruiting and initiating ribosomes or the IRES-mediated ORF is translated more quickly than that of the Cap (which would lead to fewer ribosomes along the ORF at any given time). To rule out the latter possibility, we estimated Cap-dependent and IRES-mediated elongation rates by measuring the time it takes ribosomes to run-off the biosensor after translation initiation is blocked by Harringtonine.^40^ Consistent with ribosomal run-off, Harringtonine led to a steady decay in the nascent chain signals within translation sites (Figure 4 and Figure S4A-B). This decay was not due to photobleaching because it was not observed in untreated control cells (Figure S4C-D). Fitting the linear portion of the decay gave elongation rates of 1.44 ± 0.40 aa/sec for Cap-dependent translation and 1.81 ± 2.39 aa/sec for IRES-mediated translation, corresponding to run-off times of 45 min and 43 min, respectively. The similarity of these rates and run-off times demonstrates elongation is not responsible for the lower number of IRES-mediated translation sites compared to Cap-dependent translation sites.

**Figure 4:**
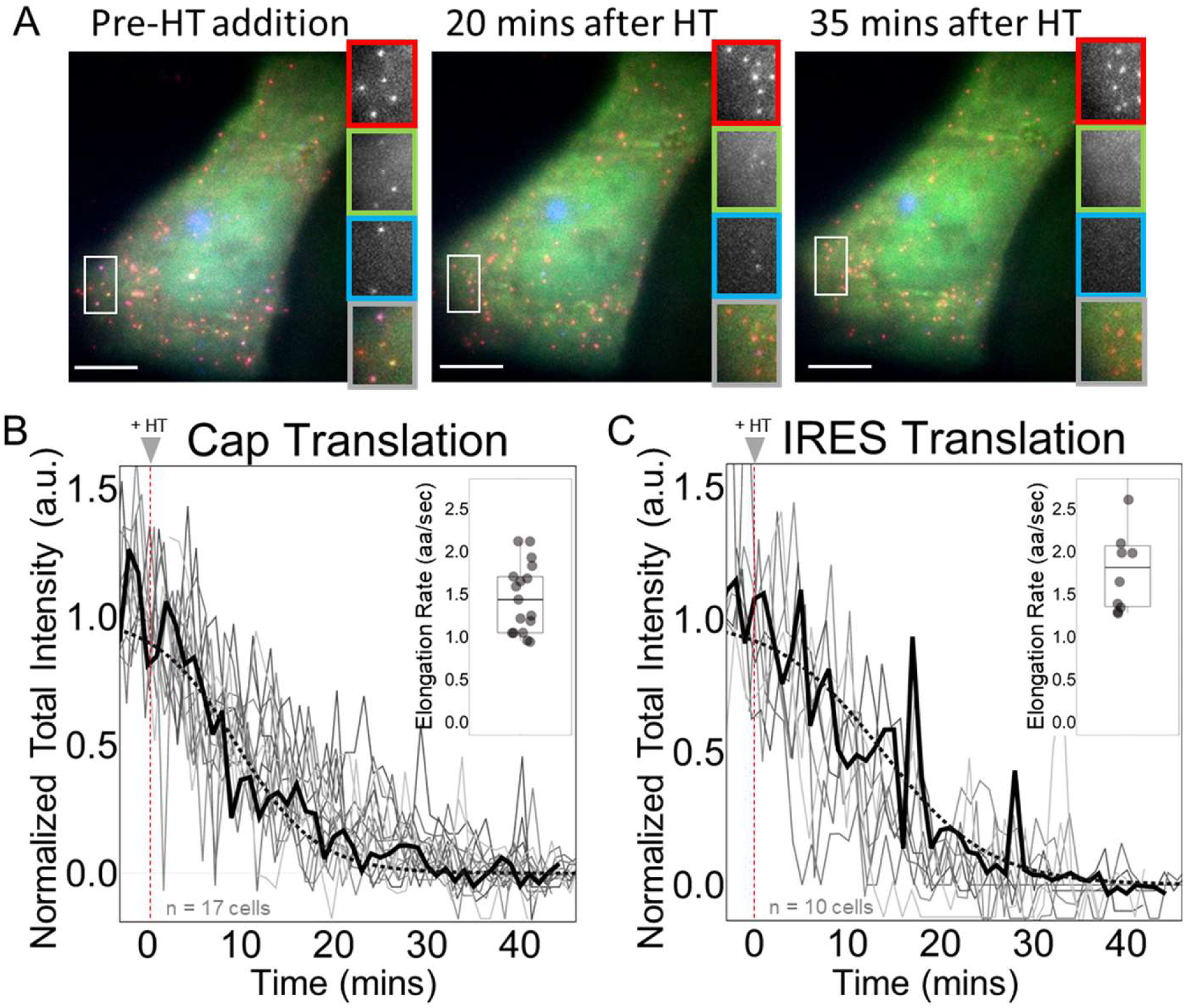
Elongation does not alter the translation efficiency of the IRES. (A) Cells before harringtonine (HT) addition, 20 minutes after HT addition, and 35 minutes after HT addition with crops of the mRNA channel (red), Cap channel (green), IRES channel (blue), and merge (gray). (B-C) Normalized total intensity of nascent chains in Cap-dependent and IRES-mediated translation sites, respectively. Each gray line represents a single cell treated with HT. The black line shows a representative cell. The dotted black line is a phenomenological fit of the representative cell. The inset is the estimated elongation rates of each cell in amino acids per second (aa/sec). All cells were imaged for 45 minutes with a 1-minute interval between each capture. Harringtonine was added after 5 captures marked by the red dotted line at time point 0. Intensity values are in arbitrary units (a.u.). n=17 cells for Cap Translation. n=10 cells for IRES Translation.

### The Cap recruits and initiates two to three times more ribosomes than the IRES

Having demonstrated elongation does not distinguish Cap and IRES translation sites, we turned our attention to ribosome recruitment and translation initiation. Assuming one of these factors limits IRES-mediated translation, we would predict fewer IRES-mediated ribosomes than Cap-dependent ribosomes in single translation sites. To test this prediction, we needed to fairly compare the intensities of nascent chain signals within single translation sites. A direct comparison was not possible because the Cap-dependent and IRES-mediated nascent chains differ in sequence, have a different number of tags, and are labeled by complementary fluorophores and probes that have different binding kinetics and different quantum efficiencies. To enable a fairer comparison, we developed a Switch Tag in which the reporters were swapped (Figure 5A). This allowed us to compare the exact same reporter under the control of both the Cap (in the Original Tag, for example) and the IRES (in the Switch Tag). In this way, we could ensure any differences in the intensity of translation sites would reflect differences in ribosome recruitment and/or initiation dynamics rather than differences in fluorophore/probe detection kinetics and/or codon biases within epitope tags.

**Figure 5:**
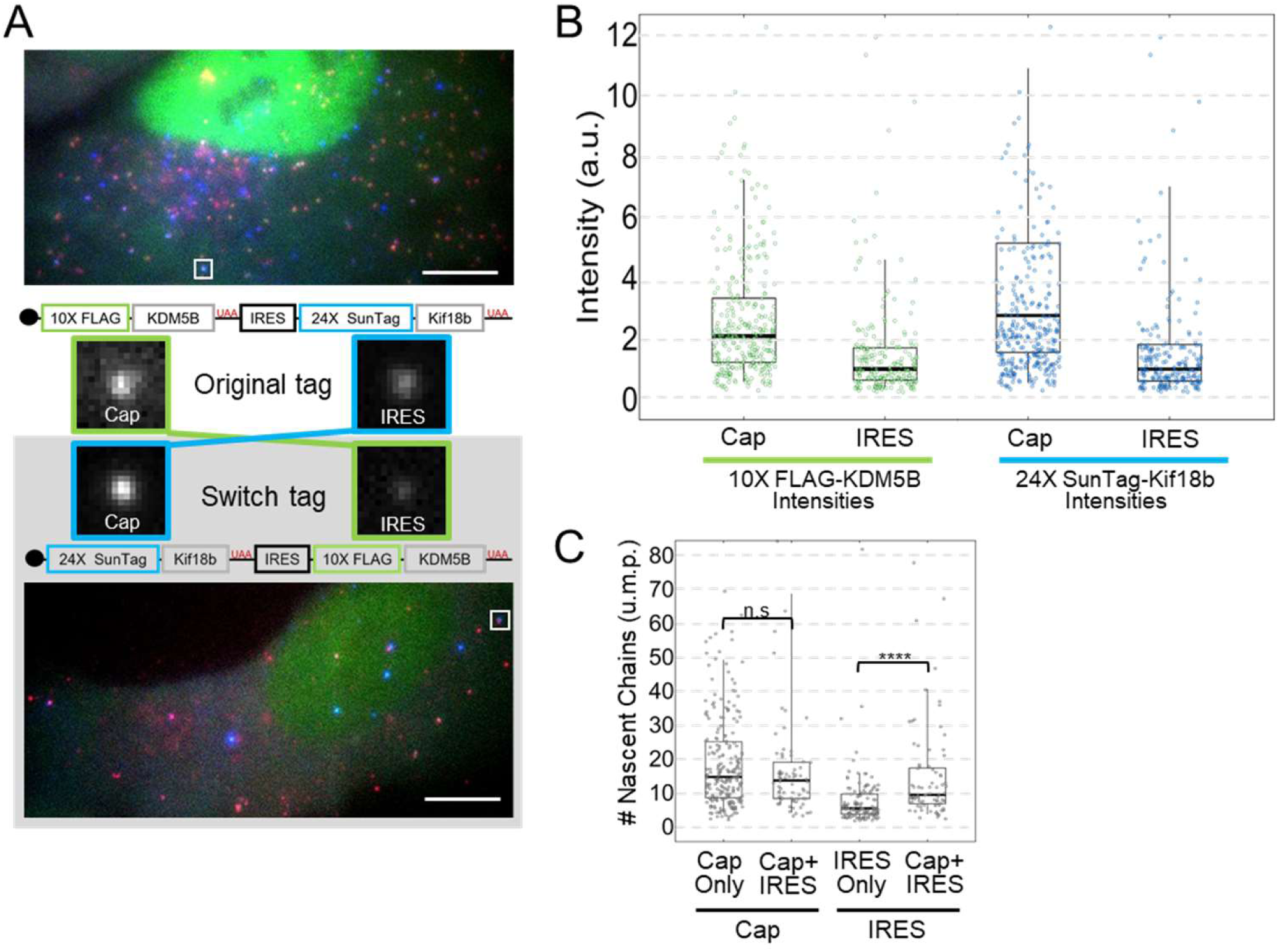
The Cap recruits and initiates two to three times more ribosomes than the IRES. (A) Representative cells from the Original Tag and Switch Tags with a representative Cap+IRES translation spot highlighted by the white square. Crops of the representative sites are shown between the cells. The construct schematic with the corresponding crop illustrates how the intensity comparisons between Cap and IRES were conducted. (B) Box and Whisker plots showing the intensity comparisons between Cap and IRES. Left graph shows intensity comparisons of 10× FLAG-KDM5B nascent chain signals from Cap in the Original Tag (n=302spots) and IRES in the Switch Tag (n=167spots). Right graph shows intensity comparisons of 24× SunTag-Kif18b from Cap in the Switch Tag (n=262spots) and IRES in the Original Tag (n=201 spots). Intensity measurements are in arbitrary units (a.u.). For the box and whisker plots, the thick black lines indicate the medians, the boxes indicate the 25%-75% range, and the whiskers indicate the 5%- 95% range. (C) Intensities of Cap in Cap Only (Original Tag) translation sites were compared to Cap in Cap+IRES (Original Tag), IRES in IRES Only (Switch Tag) and IRES in Cap+IRES (Switch Tag) to obtain numbers of ribosomes in units of mature protein (u.m.p.) on all types of translating species. Cap Only sites (n=226spots) have a median of 14.6 ribosomes, Cap+IRES sites (n=76spots) have a median of 13.6 Cap-dependent ribosomes and 9.4 IRES-mediated ribosomes. IRES Only sites (n=121spots) have a median of 5.4 ribosomes.

Reassuringly, when the Switch Tag was expressed in cells, it yielded nearly the same percentages of each type of translation site as the Original Tag (Figure S5A), demonstrating the 10×FLAG and 24×SunTag reporters do not interfere with translation dynamics and have similar detection efficiencies. As expected, there were notable differences in the intensities of translation sites. A direct comparison of the intensity of translation sites encoding 10×FLAG-KDM5B initiated in a Cap-dependent manner (from the Original Tag) versus an IRES-mediated manner (from the Switch Tag) gave a median intensity ratio of 2.1 ± 0.1 (Figure 5B). Similarly, a direct comparison of the intensity of translation sites encoding 24×SunTag-Kif18b initiated in a Cap-dependent manner (from the Switch Tag) versus an IRES-mediated manner (from the Original Tag) gave a median intensity ratio of 2.8 ± 0.2 (Figure 5B). Here we restricted our analysis to translation sites with relatively dim RNA signals to eliminate complications that could arise from co-translational mRNA clustering (Figure S5B).^27^ The similarity of the ratios we measured indicate the presence of between two and three times more ribosomes in Cap-dependent translation sites compared to IRES-mediated translation sites. In other words, for every IRES-mediated ribosome on the biosensor, there are two to three Cap-dependent ribosomes.

To extend this measurement and obtain absolute ribosome occupancies, we developed a 10×FLAG calibration construct that produces translation sites containing approximately 11 ribosomes (Figure S5C). Comparing the intensity of these translation sites to 10×FLAG-KDM5B translation sites in the Original and Switch Tags revealed that Cap-only translation sites have a median of 14.6 ribosomes (Figure 5C), while Cap+IRES translation sites have 13.6 Cap-dependent ribosomes (p value = 0.196) and 9.4 IRES-mediated ribosomes, and IRES-only translation sites have just 5.4 ribosomes (p value = 5.83E-8) (Figure 5C). Thus, Cap-dependent translation sites have more ribosomes than IRES-mediated translation sites, consistent with the higher percentage of mRNA translated in a Cap-dependent versus IRES-mediated manner. These data demonstrate that Cap-dependent translation is overall more efficient than IRES-mediated translation in our biosensor, both at the population level and single-molecule level.

### Computational modeling reveals ribosomal recruitment and not initiation limits IRES translation

Thus far, the main differences we have observed between Cap and IRES translation are (1) Cap translation occurs on a greater fraction of translation sites and (2) Cap translation sites have more ribosomes. According to our experiments, these differences are due to rate-limiting steps that precede elongation, presumably either ribosome recruitment and/or translation initiation. To distinguish these possibilities, we developed a set of theoretical models with varying levels of complexity. All models consider the kinetics of individual ribosomes translating along a single mRNA, with codon-dependent elongation proportional to the prevalence of the associated tRNA in the human genome^41^ and stochastic initiation. Models differ in the number of states an mRNA can transition between: Three-state models include an inactive mRNA state (OFF), an active mRNA state that allows Cap translation (Cap-ON), and an active mRNA that allows IRES translation (IRES-ON). Four-state models include an additional active mRNA state (Cap+IRES-ON) that allows both Cap and IRES-translation (Figure 6A).

**Figure 6:**
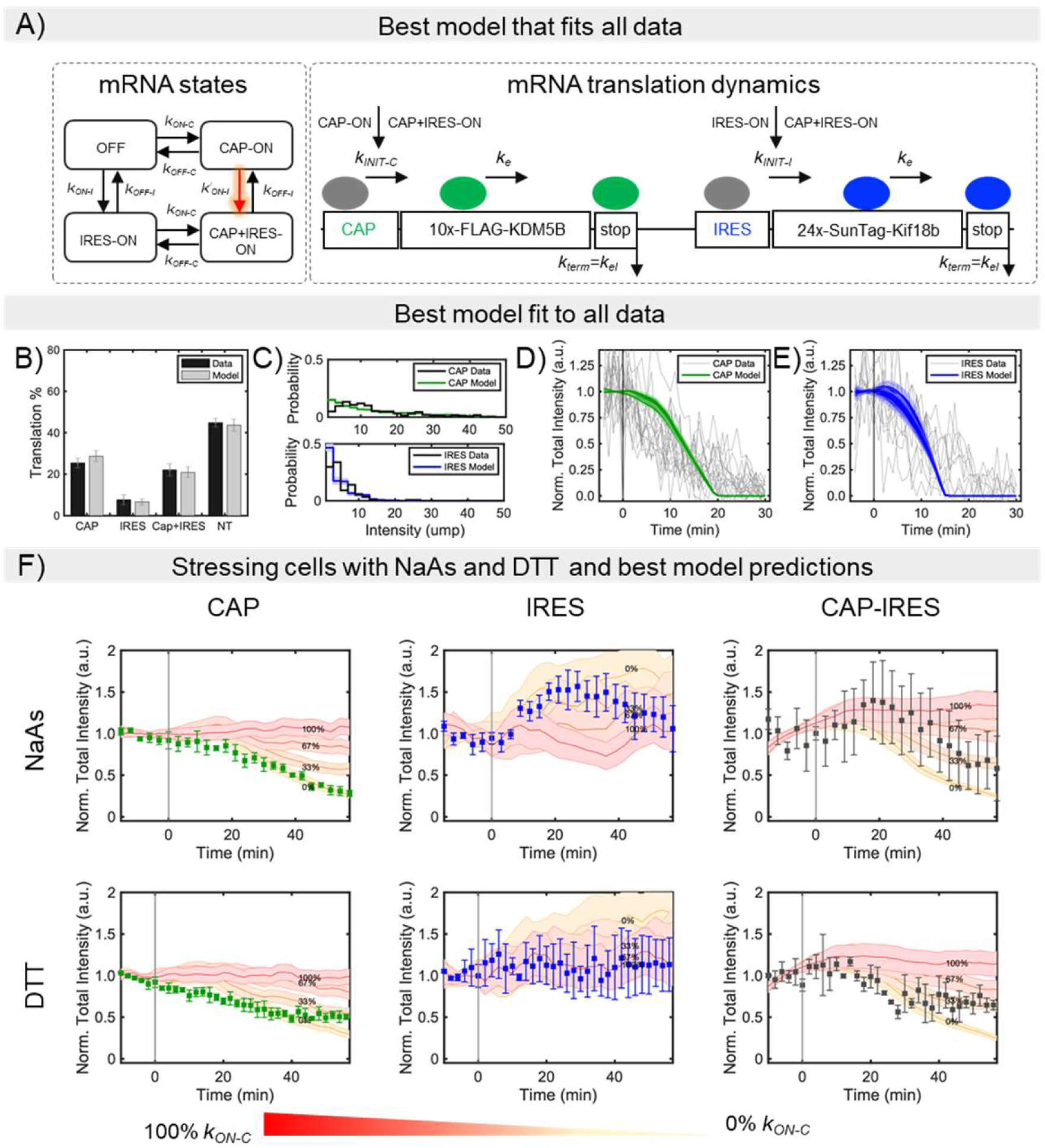
Modeling bicistronic translation of the multicolor biosensor. (A) The mathematical model considers four mutually exclusive RNA states: non-translating (OFF), Cap-dependent (CAP-ON), IRES-mediated (IRES-ON), and both Cap and IRES (CAP+IRES-ON). When in the appropriate state, initiation can take place for Cap or IRES. Elongation and termination processes continue independent of RNA state. To capture interdependence between Cap and IRES states, multiple hypotheses were tested, and the best model was selected after parameter optimization and model reduction (Figure S6C-D). The selected model considers 4 promoter states, where the IRES activation rate depends on the Cap-state (model *4S_Im2_*). (B) Model and data for the fraction of Cap-dependent (Cap), IRES-mediated (IRES), both Cap-dependent and IRES-mediated (Cap+IRES), and non-translating spots (NT). The prevalence of translation events are shown as the percentage of total RNA. (C) Experimental and model intensity distributions for Cap-dependent and IRES-mediated translation. Distributions consider only those spots that have intensities greater than or equal to one unit of mature protein (u.m.p.). (D) Decrease in intensity after Harringtonine application for Cap-mediated translation spots and (E) IRES-mediated translation spots. To denote variability, 10 independent model simulations are plotted in D and E. (F) Experimental data and simulated predictions for translation inhibition by the chemical stresses NaAs and DTT. Chemical stresses were simulated by reducing the Cap activation rates at the RNA state level (i.e. blocking *k_ON-C_*). Experimental data is represented by the square symbols. Errors bars in the experimental data and simulations are SEM. The values given in the figure represents the percentage of inhibition. Cap intensities represent Cap-only spots; IRES intensities represent IRES-only spots; and Cap+IRES intensities represent the Cap translation intensity in spots with both Cap and IRES intensities.

The stochastic dynamics for each model were simulated over large ranges of potential parameters, and we used automated parameter optimization to find combinations of mechanisms and parameters to maximize the likelihood of all quantitative data for both Cap- and IRES- translation, including the fraction of translating spots (Figure 1), Harringtonine run-off kinetics (Figure 4), and the translation site intensity distributions (Figure 5; also see Computational Methods Section and Figure 6B-E). In total, we considered 14 unique models with between 7 and 12 free parameters each, some of which included interdependence between Cap- and IRES-translation, either in the form of enhanced transition rates between states or via reinitiation of ribosomes from Cap to IRES (Equations 1 and 2; See Computational Methods Section and Figure S6A-B). Of these, the final model with minimal parameters that reproduces all experimental data has just eight parameters (Figure 6A and Table 1): (1) a baseline elongation rate of = 1.7 aa/sec, agreeing with our earlier estimate and also in line with the spectrum of elongation rates reported in the literature;^44, 45^ (2) an initiation rate *kINIT-C* ∼ 1/21 sec^-1^ for Cap-dependent translation; (3) an initiation rate *kINIT-I* ∼ 1/20 sec^-1^ for IRES-mediated translation; (4) Cap activation bursts with refractory periods (*1/kON-C*) of 34.5 min and (5) durations of (*1/kOFF-C*) of 8.3 min, leading to the synthesis of *kINIT-C /kOFF-C* = 24 nascent proteins on average per Cap burst; (6) In the absence of Cap, the model predicts that typical bursts of IRES translation would have a refractory period (*1/kON-I*) of 91.3 min and (7) a duration of (*1/kOFF-I*) of 2.5 min, leading to the synthesis of 7.5 nascent proteins on average per IRES burst. According to these fitted parameters, the efficiency of IRES translation is not limited by initiation (since *kINIT-I* ∼ *kINIT-C* ∼ 1/20 sec^-1^), but rather the IRES spends less time in a translationally active state that can recruit ribosomes.

**Table 1.**
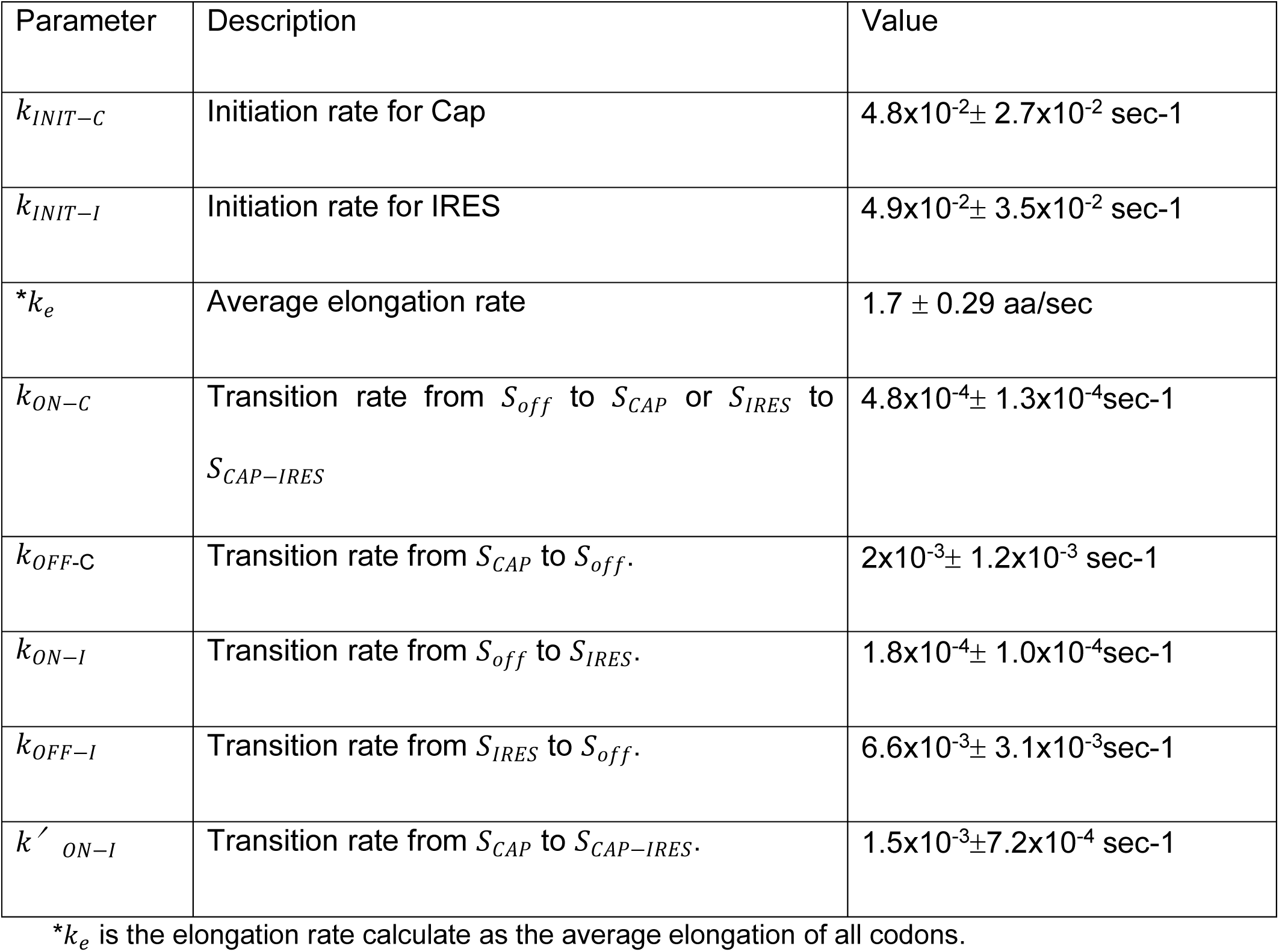
Estimated parameter values for the final selected model. (4 promoter states and promoter activation for IRES influenced by Cap).

| Parameter | Description | Value |
| --- | --- | --- |
| $k_{INIT-C}$ | Initiation rate for Cap | $4.8 \times 10^{-2} \pm 2.7 \times 10^{-2} \text{ sec}^{-1}$ |
| $k_{INIT-I}$ | Initiation rate for IRES | $4.9 \times 10^{-2} \pm 3.5 \times 10^{-2} \text{ sec}^{-1}$ |
| $*k_e$ | Average elongation rate | $1.7 \pm 0.29 \text{ aa/sec}$ |
| $k_{ON-C}$ | Transition rate from $S_{off}$ to $S_{CAP}$ or $S_{IRES}$ to $S_{CAP-IRES}$ | $4.8 \times 10^{-4} \pm 1.3 \times 10^{-4} \text{ sec}^{-1}$ |
| $k_{OFF-C}$ | Transition rate from $S_{CAP}$ to $S_{off}$. | $2 \times 10^{-3} \pm 1.2 \times 10^{-3} \text{ sec}^{-1}$ |
| $k_{ON-I}$ | Transition rate from $S_{off}$ to $S_{IRES}$. | $1.8 \times 10^{-4} \pm 1.0 \times 10^{-4} \text{ sec}^{-1}$ |
| $k_{OFF-I}$ | Transition rate from $S_{IRES}$ to $S_{off}$. | $6.6 \times 10^{-3} \pm 3.1 \times 10^{-3} \text{ sec}^{-1}$ |
| $k'_{ON-I}$ | Transition rate from $S_{CAP}$ to $S_{CAP-IRES}$. | $1.5 \times 10^{-3} \pm 7.2 \times 10^{-4} \text{ sec}^{-1}$ |

$*k_e$ is the elongation rate calculate as the average elongation of all codons.

In addition to the above seven parameters (six if we set the Cap and IRES initiation rates equal), one additional parameter was required to fit the data: an enhancement in IRES activation when Cap translation is simultaneously on (i.e. *k’ON-I* > *kON-I*). Specifically, in the presence of Cap, the IRES refractory period is reduced from 91.3 min down to 11 min, leading to a 6.9 fold increase in IRES translation. This enhancement was required to capture the large percentage of Cap+IRES translation sites (which is greater than one would predict if Cap and IRES translation were independent) (Figure 1D, left) as well as the larger number of IRES-mediated ribosomes in Cap+IRES translation sites compared to IRES-only translation sites (Figure 5C). These data and the best-fit model therefore provide evidence that translation of an upstream ORF can positively impact translation of a non-overlapping downstream ORF.

### Predicting Cap and IRES translation in response to specific cellular stresses

It is well known that viral infections cause increased levels of cellular stress. During these stressful conditions, viral RNAs continue to be translated while canonical translation decreases globally.^46^ This is thought to occur in part through IRES-mediated ribosomal recruitment to viral RNAs. Extensive studies have shown that both viral and some endogenous IRES sequences remain translationally active during certain types of cellular stress.^15, 17, 18^

To visualize the impact of stress on Cap-dependent and IRES-mediated translation at the single-molecule level, we exposed cells expressing our biosensor to NaAs^42^, which induces oxidative stress, and DTT^43^, which induces ER stress. In both cases, the intensity of Cap-only translation sites decreased significantly upon stress (Figure 6F, left). In contrast, the intensity of IRES-only translation sites remained steady or increased (Figure 6F, middle), while Cap+IRES translation sites displayed an intermediate response (Figure 6F, right). These data demonstrate that IRES-mediated translation is generally more robust in response to cellular stress than Cap-dependent translation, as would be necessary for efficient viral replication in cells during infection.

Using our best-fit model, we next attempted to predict the stress-response data. We hypothesized stress could decrease Cap-dependent translation in one of two possible ways. On the one hand, translation could be inhibited at the level of ribosome recruitment by blocking mRNA activation, i.e. by decreasing *kON-C* from its predicted equilibrium value down to zero (Figure 6F). On the other hand, translation could be inhibited at the level of initiation, i.e. by decreasing *kINIT-C* from its predicted equilibrium value down to zero (Figure S6E). We tested each mechanism under assumptions of 33%, 67% and 100% reductions to the corresponding rate (Figure S6C-D). According to the resulting simulations, the first hypothesis best predicted both the NaAs and DTT data. NaAs stress is best predicted by blocking 100% of Cap activation (log-likelihood of 184 ± 20 vs 581.8 ± 2.16 for the model in which Cap-initiation is blocked), while DTT stress is best predicted by blocking ∼67% of Cap activation (likelihood of 901.3 ± 30 vs 1022.4 ± 28 for the model in which Cap-initiation is blocked). Thus, according to these results, both oxidative and ER stress mainly shut down translation by preventing Cap-dependent mRNA activation as opposed to the direct inhibition of more downstream Cap-dependent initiation.

## DISCUSSION

The ability to track the translation of single mRNA using Nascent Chain Tracking (NCT) makes it possible to directly visualize the heterogeneity of protein expression within cells from one mRNA to another, which would not be possible with standard protein expression assays based on the detection of GFP or luciferase. Thus far, NCT has been used to investigate translation that is initiated in a Cap-dependent manner, the predominant form of translation initiation used by eukaryotes. Here we extend NCT to simultaneously investigate IRES-mediated translation, a mode of translation exploited almost exclusively by viruses to hijack host translation machinery and efficiently replicate in infected cells. By creating a single-molecule bicistronic biosensor that lights up in different colors depending on whether translation is initiated at the Cap or at the IRES, we quantify precisely when, where, and to what degree IRES sequences hijack ribosomes within living human cells.

One of the hallmarks of IRES-mediated translation is that it depends on only a subset of host translation factors.^28^ Due to the lax initiation factor requirements of IRES-mediated translation compared to Cap-dependent translation, one would think that translation of the IRES could occur in special microenvironments that, for example, are enriched or depleted in specific translation factors. In contrast to this notion, however, we find little evidence for specialized microenvironments that support IRES translation. Instead, our data suggest IRES-mediated translation sites are biophysically difficult to distinguish from Cap-dependent sites, having roughly the same translation initiation and elongation rates, similar mobilities, sizes, and spatial distributions within cells, and similar propensities to cluster near other translation sites. This overall similarity may have evolved to allow the EMCV IRES to effectively compete with the Cap for host ribosomes during infection.

According to our results, the EMCV IRES is not as efficient as the Cap mainly because it spends less time in a state that is conducive to ribosome recruitment. Our observation that burst frequencies are modulated to control translational output is reminiscent of common burst control mechanisms of transcription,^44^ and this general principle of regulation could be the natural result of sharing a common subset of initiation and elongation factors. Indeed, according to our best-fit model, bursts of IRES translation are both shorter in duration (2.5 min for IRES versus 8.3 min for Cap) and separated by longer periods of inactivity (91.3 min for IRES versus 34.5 min for Cap) than bursts of Cap translation. Given the complex secondary and tertiary structure of the EMCV IRES,^8^ which presumably undergoes dynamic conformational changes in living cells, our results suggest the IRES has trouble adopting and maintaining a conformation that can recruit ribosomes and maintain active translation. In contrast, the Cap relies on a larger set of factors, including the cap-binding protein eIF4E and scaffolding protein eIF4G. Presumably these additional factors work together to better maintain a conformation that is attractive to ribosomes and more amenable to continuous translation.

One of the most interesting observations with our biosensor was that Cap translation actually enhances that of the IRES, but not the other way around. While surprising, this does make sense in hindsight given the subset of factors the IRES requires compared to the Cap. In particular, when Cap translation is on, all factors necessary for IRES translation are present at high concentrations. The presence of all these factors nearby would enhance the overall probability the IRES gets translated. In contrast, when the IRES is on, although many translation factors are present, not all factors required for Cap translation are available, including eIF4E and eIF4G. Without these factors, Cap-dependent translation is not significantly enhanced.

The precise molecular mechanisms that govern the enhancement of IRES-mediated translation in the presence of Cap-dependent translation remain unclear. An interesting possibility given our live-cell confirmation that ribosomes generally stretch out translation sites is that the stretching somehow alters the accessibility or structure of the IRES. This could impact the IRES in a number of different ways. For example, the IRES could be stabilized (increased *kOFF* in four-state model 4Sim1), its folding could become faster (increased *kON* in four-state model 4Sim2), or possibly ribosomes coming off the Cap could reinitiate at the IRES with higher probability (the addition of *kCO*). According to our simulations, all of these possibilities do indeed improve the overall fits to our data, but faster folding alone was sufficient to improve the fit to near optimum values (see Figure S6C). Thus, our data suggest the stretching out of actively translated transcripts can impact the translation of downstream ORFs, even when those downstream ORFs do not overlap with the upstream ORF.

Despite the lower overall translation efficiency of the EMCV IRES compared to the Cap, the upside of relying on a subset of factors is IRES-mediated translation can persist and actually surpass Cap-dependent translation during stressful conditions, a situation viruses have evolved and exploited in their ongoing arms race with eukaryotic cells. We in fact see that in NaAs stress, IRES-mediated translation remains strong, presumably because this stress specifically targets eIF4E, one of the cap-binding proteins,^12^ which is not required for IRES translation. Though IRES translation also remained strong compared to Cap in DTT stress, the effect was smaller than with NaAs, presumably because DTT stress impacts a different set of translation factors than NaAs. In the future, it will be interesting to investigate which factors specifically play the biggest roles and also which IRES sequences are most robust to each type of stress. As many viruses have evolved different IRES architectures to capitalize on this distinction, there are likely an abundance of mechanisms involved in each stress, each dependent on a different set of factors with various relative timings.

Overall, our technology to visualize when, where, and to what degree IRES-mediated translation occurs at the single-molecule level in the natural setting of living cells provides a new angle on viral translation that will complement technologies like ribosome profiling^45^ and *in vitro* single-molecule assays.^46^ As quantitative single-molecule experiments and rigorous computational analyses continue to improve, we anticipate that integrated biosensors and stochastic models like those introduced here will provide new insights into how viruses exploit IRES elements and could eventually be combined with computer-aided high-throughput screens to design and verify new compounds to fight viral infection in a targeted manner.

## ACKNOWLEDGMENTS

We thank Dr. Luke Lavis for kindly providing JF646 labeled HaloTag ligand and Dr. Hataichanok (Mam) Scherman for purifying Halo-MCP and GFP-scFv (anti-SunTag). We thank all members of the Stasevich and Munsky labs for their support and helpful discussions. BM and LA were supported by a grant from the W.M. Keck Foundation and by the NIH (grant no. 5R35GM124747). TJS, AK, and TM were supported by the NIH (grant no. R35GM119728).

## AUTHOR CONTRIBUTIONS

A.K. and T.J.S. designed and planned all experiments. A.K. cloned all plasmids and performed all experiments. T.M. assisted A.K. with microscopy and particle tracking. B.M. and L.A. performed all modeling and fitting. T.J.S. and A.K. wrote the main manuscript, with assistance from T.M. for experimental methods related to microscopy and particle tracking and from B.M. and L.A. for computational sections. B.M. and L.A. wrote the computational methods. A.K., L.A., T.M., B.M. and T.J.S. edited the manuscript. B.M. and T.J.S. acquired funding and designed the computational and experimental studies.

## MATERIALS AND METHODS

### Plasmid construction

The Original Tag (SM-KDM5B-EMCV-SunTag-Kif18b-MS2) contains a spaghetti monster (SM) with 10× FLAG epitopes, a SunTag with 24× SunTag epitopes, and an MS2 repeat with 24× MS2 stem loops. The coding region of the SunTag and Kif18b was obtained by polymerase chain reaction (PCR) of a pCMV-SunTag-Kif18b-PP7 template (Addgene #128606), using the following primers: 5’-GCC GAA AGG TTT AAA CGC TAG CTC TGG AGG AGA AGA ACT TTT GAG CAA GAA T-3’; 5’-AGT AAC AGT CCG CCT AGG TCC TTA TCG GAC ACC TTG GT-3’. The PCR product contained arms of homology to the acceptor plasmid (SM-KDM5B-MS2; Addgene #81084). The acceptor plasmid was cut with NheI (New England BioLabs) between the end of KDM5B and the MS2 stem loops. The PCR product and cut acceptor plasmid were assembled via Gibson Assembly (homemade mixture). The resulting plasmid was SM-KDM5B-Nhe1-SunTag-Kif18b-MS2, which was also used as the NoIRES construct. The EMCV IRES sequence was amplified by PCR from EMCV_IRES_pcDNA4TO_H2B_SunTag24x_v1 (Addgene #246719) using the following primers: 5’-CCG AAA GGT TTA AAC GCT AGC ACG TTA CTG GCC GAA-3’; 5’-TTC TTC TCC TCC AGA GCT AGC TAT TAT CAT CGT GTT TTT CAA AGG AAA-3’. The PCR product contained arms of homology to the acceptor plasmid (SM-KDM5B-Nhe1-SunTag-Kif18b-MS2). The acceptor plasmid was cut with NheI (New England BioLabs) between the end of KDM5B and beginning of SunTag. The PCR product and cut acceptor plasmid were assembled via Gibson Assembly. The start codon for SunTag-Kif18b is within the EMCV IRES sequence.

For the construction of the Switch Tag (SunTag-Kif18b-EMCV-SM-KDM5B-MS2), the coding region of the SunTag and Kif18b was obtained by PCR of a pCMV-SunTag-Kif18b-PP7 template (Addgene #128606), using the following primers: 5’-TCG CTG TGA TCG TCA CTT GGC GGA CAC CAT GGA AGA ACT TTT GAG CAA GAAT-3’; 5’-CGT CCT TGT AGT CCA TGG TGG CGG CGC GCC GTC TTA GAT ATC GGA CAC CTTG-3’. The PCR product contained arms of homology to the acceptor plasmid (SM-KDM5B-MS2 Addgene #81084). The acceptor plasmid was cut with NotI (New England BioLabs) at the beginning of SM. The PCR product and cut plasmid were assembled via Gibson Assembly. The resulting plasmid was SunTag-Kif18b-Nhe1-SM-KDM5B-MS2. The EMCV IRES sequence was amplified by PCR from EMCV_IRES_pcDNA4TO_H2B_SunTag24×_v1 (Addgene #246719) using the following primers: 5’-CCA AGG TGT CCG ATA TCT AAG ACG GCG TTA CTG GCC GAA GCC GCT-‘3; 5’-CCT TGT AGT CCA TGG TGG CGG CGC ATA TTA TCA TCG TGT TTT TCA AAG GAA AAC CAC-3’. The PCR product contained arms of homology to the acceptor plasmid (SunTag-Kif18b-AscI-SM-KDM5B-MS2). The acceptor plasmid was cut with AscI (New England BioLabs) between the end of Kif18b and beginning of SM. The PCR product and cut acceptor plasmid were assembled via Gibson Assembly. The start codon for SM-KDM5B is within the EMCV IRES sequence.

### anti-FLAG Fab generation and dye-conjugation

Pierce mouse IgG1 preparation kit (Thermo Scientific) was used to generate Fab according to the manufacturer’s instruction. Briefly, immobilized ficin in the presence of 25 mM cysteine was used to digest FLAG antibodies (Wako, 012-22384 Anti DYKDDDDK mouse IgG2b monoclonal) to create Fab. Fab were separated from the Fc region using NAb Protein A column. After elution, Fab were concentrated to 1 mg/ml and conjugated to Cy3. Cy3 N-hydroxysuccinimide ester (Invitrogen) was dissolved in DMSO and stored at −20°C. 100 µg of Fab were diluted into 100 µl of 100 mM NaHCO3 (pH 8.5). 1.33 µl of Cy3 was added to this solution and incubated with end-over-end rotation for 1-2 hours at room temperature. The conjugated Fab were then eluted from a PBS pre-equilibrated PD-mini G-25 desalting column (GE Healthcare) that removed unconjugated dye. Conjugated Fabs were then concentrated using an Ultrafree 0.5 filter (10k-cut off; Millipore) to 1 mg/ml. The Fab:dye ratio was calculated using the absorbance at 280 and 550 nm, and using the extinction coefficient of Fab with the dye correction factor at 280 nm provided by the manufacturers (0.08 for Cy3). The degree of labeling was calculated using the following formula:

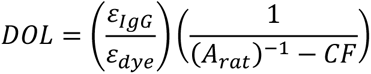

### MCP and scFv-GFP purification

His-tagged MCP/scFv-GFP was purified with Ni-NTA-agarose (Qiagen) following the manufacturer’s instructions with minor modifications. Briefly, the bacteria were lysed in a PBS-based buffer containing a complete set of protease inhibitors (Roche), binding to the Ni-NTA resin was carried out in the presence of 10 mM imidazole. After washing with 20 and 50 mM imidazole in PBS, the protein was eluted with 300 mM imidazole in PBS, and directly used for experiments. The rest was dialyzed against a HEPES-based buffer (10% glycerol, 25 mM HEPES pH 7.9, 12.5 mM MgCl2, 100 mM KCl, 0.1 mM EDTA, 0.01 % NP-40 detergent, and 1 mM DTT) and stored at −80 °C after snap-freezing by liquid nitrogen.

### Cell culture, transfection, and beadloading

U2OS cells were grown in DMEM (Thermo Scientific) supplemented with 10% (v/v) FBS, 1 mM L-glutamine and 1% (v/v) Penicillin-streptomycin (DMEM+). One to two days prior to experiments, cells were plated into a 35 mm MatTek chamber at approximately 70-80% confluency. Two to four hours prior to experiments, cells were put in OPTI-MEM (Thermo Scientific) supplemented with 10% (v/v) FBS (OPTI-MEM+). Cells were then bead-loaded with fluorescently labeled Fab, GFP-fused scFv, MCP-HaloTag protein and purified DNA of interest. Briefly, 100 µg/ml of fluorescently labeled Fab, 100 µg/ml of purified GFP-fused scFv, 33 µg/ml of purified MCP HaloTag protein, and 750ng of DNA of interest were prepared in a total volume of 4µl of 1xPBS. After removing OPTI-MEM and FBS, the 4µL solution was pipetted to the top of the cells. Then, ∼106 µm glass beads (Sigma Aldrich) were sprinkled evenly over the cells. The chamber was tapped firmly 12 times on the bench, and OPTI-MEM+ was added back to the cells. Two hours after bead-loading, cells were washed twice with phenol-red-free DMEM+ such that all beads were removed. 200 nM of JF646-HaloTag ligand was next added (1µL of 200nM to 1mL of phenol-red-free DMEM+). After 20 minutes of incubation at 37 ֯ C, the cells were washed twice with phenol-red-free DMEM+ to remove excess ligand. 2 mL of phenol-red-free DMEM+ was added back to the cells. Translation experiments were conducted immediately after washing.

### Single molecule tracking microscopy

To track single molecule mRNA translation events, we used a custom-built widefield fluorescence microscope with a highly inclined illumination scheme.^24, 31^ Briefly, the excitation beams, 488, 561 and 637 nm solid-state lasers (Vortran), were coupled and focused on the back focal plane of the objective (60X, NA 1.49 oil immersion objective, Olympus). The emission signals were split by an imaging grade, ultra-flat dichroic mirror (T660lpxr, Chroma) and detected by two aligned EM-CCD cameras (iXon Ultra 888, Andor) by focusing with 300 mm tube lenses (this lens combination produces 100X images with 130 nm/pixel). Live cells were placed into an incubation chamber (Okolab) at 37 °C and 5% CO2 on a piezoelectric stage (PZU-2150, Applied Scientific Instrumentation). The focus was maintained with the CRISP Autofocus System (CRISP-890, Applied Scientific Instrumentation). Image acquisition was performed using open source Micro-Manager.^47^ With this setting, one camera detected far-red emission signals while the other detected either red or green emission signals.

Far-red signals were excited with the 637 nm laser with a 731/137 nm emission filter (FF01-731/137/25, Semrock). Red and green signals were separated by the combination of the excitation lasers and the emission filters installed in a filter wheel (HS-625 HSFW TTL, Finger Lakes Instrumentation); namely, the 561 nm laser and 593/46 nm emission filter (FF01-593/46-25, Semrock) were used for Cy3 imaging, and the 488 nm laser and 510/ 42 nm emission filter (FF01-510/42-25, Semrock) were used for sfGFP or A488 imaging. The lasers, filter wheel, cameras, and the piezoelectric stage were synchronized by an Arduino Mega board (Arduino). The exposure time of the cameras was selected as 53.64 msec throughout the experiments. The readout time for the cameras from the combination of imaging size, readout mode, and the vertical shift speed was 23.36 msec, resulting in an imaging rate of 13 Hz (77 msec per image). The excitation laser lines were digitally synched to ensure they only illuminated cells when the camera was exposing to avoid excessive photobleaching. To capture the entire volume of the cytoplasm of U2OS cells, 13 z stacks with a step size of 500 nm (6 μm in total) were acquired using the piezoelectric stage. Because one image of Cy3 was captured on one camera and one image of sfGFP/A488 + JF646 was captured on the other camera in the same stack of the cell, the z-position within the cell changed every two images. The position of the filter wheel was changed during the camera readout time. This resulted in a total cellular imaging rate of 0.5 Hz (2 s per volume for 3-colors). Note that all colors described in the text and that are shown in the figures are based on the color of the excitation laser: RNA in red (JF646) and protein in green (Cy3) or blue (sfGFP).

### Cell imaging conditions with no drugs added for all constructs

Cells beadloaded with SM-KDM5B-EMCV-SunTag-Kif18b-MS2 (Original Tag), SunTag-Kif18b-EMCV-SM-KDM5B-MS2 (Switch Tag), or SM-KDM5B-SunTag-Kif18b-MS2 (NoIRES Tag), Cy3 labeled anti-FLAG Fab, Halo-MCP protein (labeled with JF646-HaloTag ligand), and anti-SunTag scFv-GFP were imaged with a 6 second interval between every 13 captures (one entire cell volume) for 25-50 total time-points. Laser powers for all images were: 15mW for 637nm, 9mW for 488nm, and 5mW for 561nm with an ND10 neutral density filter at the beam expander.

### Particle tracking

Collected images were first pre-processed with Fiji.^48^ Briefly, the 3D images were projected to 2D images by a maximum intensity projection and background subtracted. Post-processed images were then analyzed by a custom-written *Mathematica* (Wolfram Research) routine to detect and track particles in the RNA channel (red color). Specifically, particles were emphasized with a band-pass filter such that the positions could be detected using the built-in *Mathematica* routine ComponentMeasurements ‘‘IntensityCentroid.’’ Detected particles were linked through time by allowing a maximum displacement of 5 pixels between consecutive frames. Particle tracks lasting at least 5 frames were selected. To properly account for the offset between the two cameras, a geometric transformation function (see method below) was applied to the coordinates of the center of mRNAs. For each frame of each track, 15×15 (pixels × pixels) crops centered on the registered mRNA coordinate were made and averaged through time.

Using *Mathematica’s* bandpass filter and ComponentMeasurements described above, the time-averaged crops corresponding to each track were categorized based on the presence of detectable signals in the green and blue nascent chain channels: Red – mRNA not translating, Red + Green = Yellow – mRNA translating in Cap Only for Original Tag or IRES Only for Switch Tag, Red + Blue = Purple – mRNA translating in IRES Only for Original Tag and Cap Only for the Switch Tag, Red + Green + Blue = White – mRNA translating in both Cap and IRES manner. Once the spots were categorized in this automated fashion, all spots were again hand-checked to minimize error.

Finally, the original 2D maximum intensity projected images corresponding to each hand-checked track were fit to find their precise coordinates and intensities (using the built-in *Mathematica* routine NonlinearModelFit) to a 2D Gaussians of the following form:

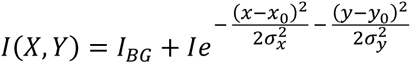

where *I_BG_* is the background intensity, *I* the particle peak intensity, (**σ*_x_, *σ*_y_*) the spread of the particle, and (*x*_0_, *y*_0_) the particle location. From these data, the intensity, position through time, and number of spots over time in each track were quantified for downstream analysis.

### Fast imaging for mean square displacement analysis

For fast particle tracking to accurately quantify the mean square displacements, single planes of cells loaded with the Original Tag construct, Cy3 anti-FLAG Fab, anti-GCN4 scFv-GFP, and Halo-MCP were imaged with an imaging rate of 77 msec.

### Geometric transformation function

The offset between the two cameras was registered using the built-in *Mathematica* routine FindGeometricTransform. To find the transformation function that best aligned the fitted positions, 100 nm diameter Tetraspeck beads evenly spread out across the image field-of-view were imaged on the same day experiments were taken. Only the fitted particle positions were registered to avoid introducing any distortion into images. Therefore, a slight offset can be observed between the red and the green/blue particles even though they are within a diffraction limited spot according to our registration.

### Calibrating translation site intensity

To calibrate the nascent chain intensity signals within translation sites to units of mature protein (i.e., to units that are roughly equivalent to the number of nascent chains or ribosomes within the translation site), two calibration constructs were imaged.^24^ The two calibration constructs were equal in length, one containing the spaghetti monster 10×FLAG tag (SM-BetaActin) used in the Original and Switch Tag biosensors, the other containing just a single FLAG tag (1×FLAG-filler-BetaActin). The latter was used to measure the number of ribosomes translating in a translation site. With the 1×FLAG-filler-BetaActin calibration construct, each nascent chain in a translation site contains just one FLAG epitope labeled by a single Fab conjugated (on average) to a single Cy3 fluorophore. By imaging this 1×FLAG construct at high laser powers such that translation sites and single Cy3 fluorophores (confirmed by single-step photobleaching, see below) can both be visualized, the ratio of nascent chain signals in translation sites to single Cy3 fluorophore intensity signals approximates the number of nascent chains (or ribosomes) per translation site. From these measurements, 11.4 ± 2.0 ribosomes were estimated to be translating the 1×FLAG-filler-BetaActin calibration construct. Since the 1×FLAG-filler-BetaActin calibration construct and the SM-BetaActin calibration construct (with 10×FLAG) are the same length with the same promoters and 3’ and 5’ UTRs, their translation sites should contain roughly the same number of ribosomes. With that assumption, the SM-BetaActin calibration construct can be used as an intensity calibration for all other reporters tagged with spaghetti monsters in the Original and Switch Tags, i.e. for Cap translation sites with the Original Tag and IRES translation sites with the Switch Tag.

To perform this calibration, cells were imaged in a single plane at high laser powers (50 mW for 561nm and 15mW for 637nm laser). A short movie was acquired, after which cells were continually imaged (without acquiring a movie) to photobleach them to the point at which single probe fluorescence could easily be detected by single-step photobleaching. At this point, a second short 250-frame movie was acquired. The intensity of polysomes (verified by the presence of an RNA signal intensity) from the first frame of the first movie was then measured (as described in the ‘Particle tracking’ section above) and compared to the plateau intensity of a single probe just prior to single-step photobleaching. Calibration was performed in this way to ensure at the beginning of the movie the FLAG epitopes would be close to saturated by anti-FLAG Fab. Had a lower concentration of probe been used in the beginning (although this would enable single probe tracking without the need for photobleaching), the FLAG epitopes in translation sites would be less saturated, which would lead to an underestimate of the translation site signal intensity. From these measurements, the average number of nascent chains (or ribosomes) in a translation site can be estimated from the intensity ratio of polysomes to single probes.

After calibration, cells beadloaded with anti-FLAG Fab (Cy3) and transfected with either the 10×FLAG-BetaActin calibration construct or the Original Tag construct were imaged on the same day with the same imaging conditions (50 mW for 561nm and 15mW for 637nm laser). From these experiments, the intensity of nascent chain signals in translation sites for the 10×FLAG-BetaActin calibration construct (with a known number of ribosomes of 11.4) was directly compared to that of Cap Only translation sites using the Original Tag. According to these measurements, the number of ribosomes translating in a Cap Only manner in the Original Tag is 14.6 ± 5.6 ribosomes. Comparing the intensity of these sites to all other SM translation sites using the Original Tag (Cap in Cap+IRES translation sites) and Switch Tag (IRES in IRES Only and Cap+IRES translation sites) gave the number of ribosomes translating in all possible translation sites, as shown in Figure 5C.

### Ribosome run-off experiments using Harringtonine treatment and elongation estimates

To measure average elongation rates, cells beadloaded with the Original Tag (SM-KDM5B-EMCV-SunTag-Kif18b-MS2), Cy3 labeled anti-FLAG Fab, Halo-MCP protein (labeled with JF646-HaloTag ligand), and anti-SunTag scFv-GFP were imaged with a 60 second interval between every 13 frames (one entire cell volume) for 50 total time-points. Laser powers were the same as previously described for general imaging. After acquiring 5 time-points of pre-treated images, cells were treated with a final concentration of 3 µg/mL of Harringtonine (Cayman Chemical). After treatment, cells were imaged for the remaining 45 time-points as described. As a photobleaching control, cells were imaged at the exact same imaging conditions described previously however no drug was added.

To generate ribosomal run-off curves, images were analyzed with the particle tracker as previously described. In each frame of each cell image, nascent chain signal intensities from all translation sites were totaled resulting in an intensity decay curve over time for each individual cell. Each decay curve was normalized to the average value from the first five frames (preceding drug addition). Each individual curve was fit to the following phenomenological equation:

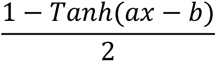

where the linear part of the fit (the slope at ax=b) provides a good estimate of the elongation rates in amino acids over time in seconds.^49^ From this, elongation rates for each cell were calculated.

### Puromycin treatment

Cells beadloaded with the Original Tag (SM-KDM5B-EMCV-SunTag-Kif18b-MS2) or the Switch Tag (SunTag-Kif18b-EMCV-SM-KDM5B-MS2), Cy3 labeled anti-FLAG Fab, Halo-MCP protein (labeled with JF646-HaloTag ligand), and scFv-GFP were imaged with a 60 second interval between every 13 frames (one entire cell volume) for 15 total time-points. After acquiring 5 time-points of pre-treated images, cells were treated with a final concentration of 50µg/mL of puromycin. After treatment, cells were imaged for the remaining 10 time-points as described previously. As a photo-bleaching control, cells were imaged at the exact same imaging conditions described previously with no drug added. Three biological replicates were taken.

### Sodium Arsenite (NaAs) and Dithiothreitol (DTT) treatment

Cells beadloaded with the Original Tag (SM-KDM5B-EMCV-SunTag-Kif18b-MS2) or Switch Tag (SunTag-Kif18b-EMCV-SM-KDM5B-MS2), Cy3 labeled anti-FLAG Fab, Halo-MCP protein (labeled with JF646-HaloTag ligand), and anti-GCN4 scFv-GFP were imaged with a 180 second interval for NaAs and 120 for DTT between every 13 frames (one entire cell volume) for 35 total time-points. After acquiring 5 time-points of pre-treated images, cells were treated with a final concentration of 0.5 mM of NaAs or 0.75 mM of DTT. After treatment, cells were imaged for the remaining time-points. As a photo-bleaching control, cells were imaged at the exact same imaging conditions described previously with no drug added. Four biological replicates were taken.

### Computational Details

A stochastic model was implemented to simulate Cap/IRES activation, ribosome initiation, elongation, termination, and potential ribosome recycling mechanisms for Cap-dependent and IRES-mediated genes.

In the mathematical model, initiation events are dictated by the mRNA state. Specifically, four possible mRNA activation states were proposed (*S_OFF_*, *S_CAP_*, *S_IRES_*, *S_CAP-IRES_*), where: *S_OFF_* represents a non-permissive initiation state; *S_CAP_* allows for only Cap-dependent ribosomal initiation; *S_IRES_* allows for only IRES-mediated initiation; and *S_CAP-IRES_* allows both Cap-dependent and IRES-mediated initiation. Eq. 1 represents the transition reactions between mRNA states.

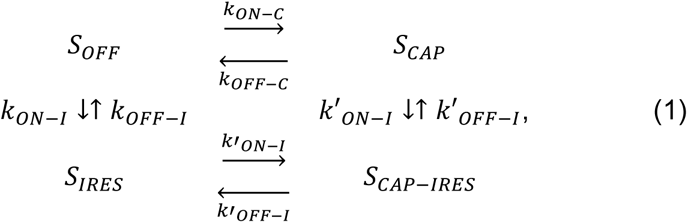

where each *k_x_* represents a first-order transition rate between two RNA states, and *k′_X_* is the transition rate conditioned on the activation state of the other construct (e.g., *k′_ON-I_* is the Cap-dependent activation rate of IRES). A simpler three-state model was considered by removing the fourth RNA state. The parameter estimation section describes how a system with three or four mRNA states was chosen.

When the system is in one of the appropriate mRNA activity states, Cap-dependent and IRES-mediated initiation events occur with propensities *w_INIT-C_* and *w_INIT-I_*, respectively, which are defined:

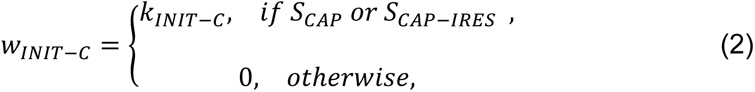

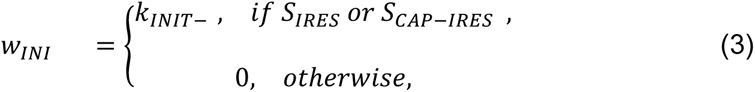

where *k_INIT-C_* and *k_INIT-_* represent the Cap and IRES initiation rates, respectively.

To simulate the model under stochastic dynamics, Eq. (2) and (3) were used to generate a vector of random initiation event times *τ_INIT_* for each gene, that is *τ_INIT_IRES__* and *τ_INIT_CAP__*. A codon-dependent model for translation was used, in which the elongation rate for each codon is given by 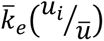, where *μ_i_* is the known frequency of the *i^th^* codon in the human genome, *k̄* is the average codon usage frequency in the human genome, and *k^-^_e_* is the basal elongation rate (to be estimated from the data). In the models, the final codon termination rates are assumed to be equal to the average elongation rate.

For increased computational efficiency, ribosome elongation is approximated using a coarse-grained procedure. For this, sparse ribosome loading were first assumed to calculate the average theoretical time needed by a ribosome to complete gene elongation, *τ_ke_*, as follows:

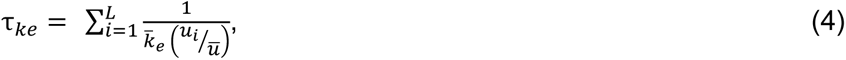

where *L* represents the gene length in codons. Using the specific gene sequence for the Cap-dependent gene and IRES-mediated gene, we calculated the total elongation time *τ_Cap_* and *τ_IRES_*, respectively. At any time, *t*, such that *0 < *t* − *τ*_INITCap_ < *τ*_Cap_*, the position of a given Cap-translating ribosome can be obtained by calculating the proportion of elongated gene as follows:

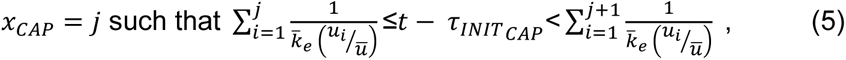

and for the IRES-mediated gene for *0 < *t* − *τ*_INIT_IRES__ < *τ*_IRES_*:

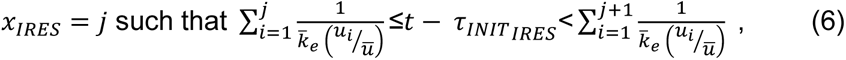

where *τ_INIT_CAP__* and *τ_INIT_IRES__* are the times at which the corresponding ribosome initiated translation begins.

To consider potential interaction mechanisms between Cap-dependent and IRES-mediated translation, two possible hypotheses were postulated:

A first hypothetical model considers potential ribosome recycling (or crossover) mechanisms, by which a ribosome that completes translation of the Cap-dependent gene could immediately re-initiate translation of the IRES-mediated gene. In this context, a new propensity, *w_CI_*, that specifies the probability that a ribosome completing Cap will re-initiate at IRES was introduced. The specification of such reactions reflects single-mRNA translation observations by Wang et al., 2016^43^, which suggest ribosome hops between adjacent open reading frames on a single RNA. To test if such recycling mechanisms are necessary to reproduce the experimental data, multiple models with and without nonzero values for the crossover rate *w_IC_* were compared.

In the second hypothetical model, Cap and IRES dependency were tested by assuming that the activation and deactivation of Cap or IRES could depend on the activity state for the other sensor (e.g., IRES could activate faster when Cap is already active). Including different combinations of these hypothetical mechanisms in the three- and four-state models led to propose a list of 14 different sub-models, each comprising between 7 and 12 free parameters (see Supplementary Figure 6-B).

### Converting ribosome elongation times to fluorescence intensity

To relate the ribosome elongation times to fluorescence intensity, a similar approach as in Aguilera et al.^41^ was adopted. Ribosome occupancy is converted to fluorescence intensity by increasing intensity after each ribosome moves across the tag-region. For this, a cumulative *probe design vector* was defined that records the number of probe sites upstream from each codon, ***c_g_*** = [*c*_1_, *c*_2_, … , *c_L_*], for the appropriate construct (i.e., *g =* Cap-dependent or IRES-mediated genes, respectively). Using this, the intensity was calculated as the sum of the product of the position of the ribosome at a given time and ***c_g_***. For Cap-dependent spots, the intensity vector is defined as:

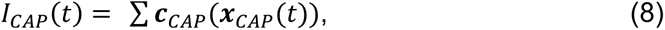

and for IRES-mediated spots it is:

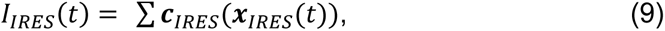

where ***c**_g_*(***x***_g_) is the intensity of a given ribosome at position ***x**_g_*, and the summations are taken over all ribosomes present on the mRNA at time *t*. To have consistent units of intensity between model simulations and experimental data, intensity values are reported in units of mature proteins (u.m.p.) as described in detail above.

### Comparison of experimental data and model

To reproduce experimental data, the model was simulated using a modified Direct Method^50^ for 4000 trajectories representing independent RNA spots. Simulations were run for a burn-in period of 10,000 seconds to approximate steady state. Simulations were processed and used to capture spot intensity for the Cap-dependent gene *(*I*_CAP_)* and the IRES-mediated gene *(*I*_IRES_)*. Additionally, simulated spots were classified as Cap-dependent with probability *P_CAP_*, IRES-mediated with probability *P_IRES_*; both with probability *P_CAP-IRES_*, or neither with probability **P*_None_*.

### Modeling Harringtonine experiments

Harringtonine inhibits new initiation events by directly blocking the 60S subunit in the ribosome, and it has been widely used to perform run-off assays to estimate elongation rates.^24^ To mimic the effects of Harringtonine in our model, the initiation rate was modified for the first gene as follows:

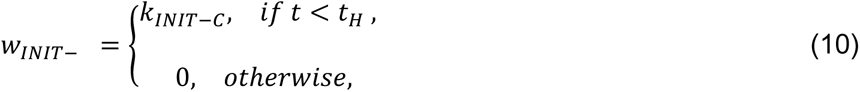

and the initiation rate for the second gene as follows:

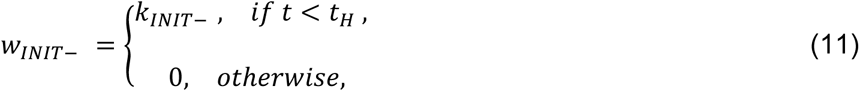

where *t_H_* is the time of application of Harringtonine.

### Modeling Sodium Arsenite (NaAs) and Dithiothreitol (DTT) *experiments*

NaAs and DTT are chemical stresses that have been used to affect Cap-dependent initiation in previous single-molecule translation experiments.^43^ The mechanism of action for NaAs is not well understood, but it has been suggested to affect ribosome initiation through its action on translation factors, such as eIF2a and eIF4.^17^ To simulate these chemical stresses, two potential mechanisms of action were tested. The first potential mechanism of action involves blocking Cap-dependent translation by affecting its RNA state, that is implemented in the model by modifying the Cap activation rates, *k_ON-C_* and *k′_ON-C_*, as follows:

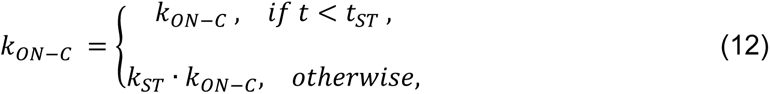

and

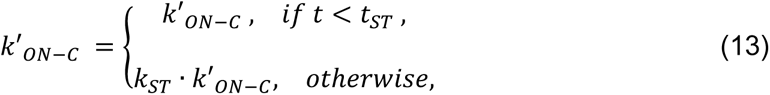

where, *k_ST_* is an inhibition constant, where a total inhibition is achieved by *k_ST_ = 0*, and a null inhibition is achieved by *k_ST_ = 1*. *t_ST_* represents the time of stress application.

In the second mechanism of action, it was hypothesize that the drug directly blocks Cap-dependent translation initiation. In the model, this is achieved by modifying *w_n_* as follows:

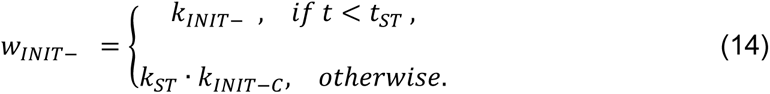

### Parameter estimation and optimization routines

The parameter estimation strategy consists of finding a parameter set (*Λ^-^*) that statistically reproduces all experimental data, including intensity histograms, fractions of translating spots, and Harringtonine ribosomal run-off assays as follows:

### Intensity histograms

To compare experimental and simulated steady-state intensity histograms, the probability to observe the experimentally determined intensities (*d*) was estimated given a parameter set (*Λ*) in the model implementation. To estimate *P(d; Λ)*, histograms were collected using *N_t_ = 4000* independent stochastic trajectories per parameter evaluation. The likelihood function was estimated as follows:

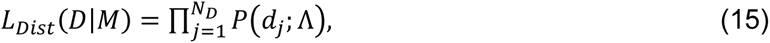

and the log-likelihood as:

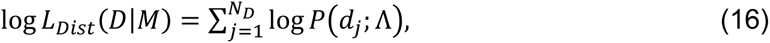

where *D* represents the data measured in *N_D_* independent experimental data and *M* corresponds to the model. As the experimental measurements can only detect protein intensities above a threshold of one mature protein, all spots with intensities below 1 U.M.P. were defined as non-translating mRNA. This metric was applied to experimental data consisting of Cap-dependent spots (CAP), and IRES-mediated spots (IRES). With this, a total log-likelihood function was calculated as the sum of the functions for Cap and IRES spots, that is:

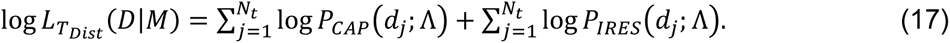

### Fraction of translating spot

A similar approach was used to compute the likelihood to observe the experimentally determined number of spots classified as Cap-only, IRES-only, Cap+IRES, and non-translating. The likelihood function was computed as follows:

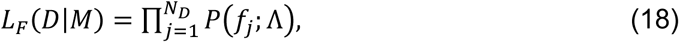

and the log-likelihood as:

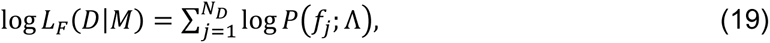

where each *f_j_* corresponds to one of the four spot types (i.e., Cap, IRES, Cap+IRES, or non-translating), *N_D_* is the number of independent observed spots, and *P(f_j_; Λ)* is the categorical distribution of spots of each type estimated by the model simulations with parameters *Λ*.

### Harringtonine induced ribosomal run-off

To compare simulated and experimental time course data representing the intensity after Harringtonine application, a Gaussian likelihood function was assumed and calculated as follows:

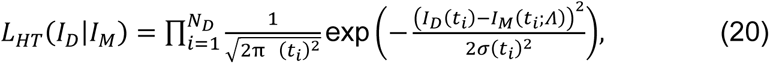

with a log-likelihood form given by:

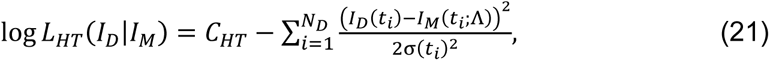

where *σ(*t*)* is approximated by the measured SEM, and *N_D_* is the number of time points from the Harrintonine run-off curve. In this log-likelihood formulation, *C_HT_* is a constant that doesn’t depend on the parameters.

Experimental data was quantified for the total intensities for Cap (*I_CAP-D_*) and IRES (*I_IRES-_*) within all spots. These two data sets were collected to compute a total log-likelihood function as follows:

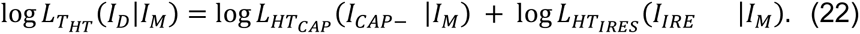

Parameter searches consisted of optimization routines based on genetic algorithms (GA) using the function *ga* in MATLAB. The optimization routine was implemented with a population of 100 individuals for 30 generations, and the implementation was run multiple times with random initial conditions. Additionally, the Pattern Search Algorithm^51^ was implemented using the function *patternsearch* in MATLAB to ensure convergence. The best parameter values were selected by minimizing a global objective function that considers all data sets, that is:

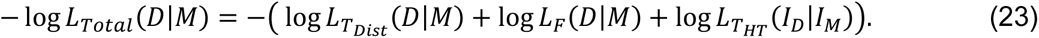

The comparison of the optimization results for all tested models is given in Supplementary Figure 6-C and D.

### Assessing how well models predict Sodium Arsenite (NaAs) and Dithiothreitol (DTT) experiments

After optimizing the models, cross-validation experiments were predicted using the chemical stresses, NaAs and DTT. For this, simulated and experimental time course data representing the total translation spot intensity after NaAs or DTT application were compared. The likelihood function was calculated as follows:

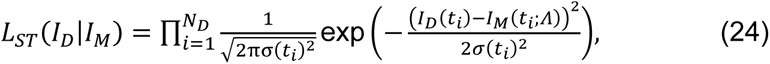

and the log-likelihood function is:

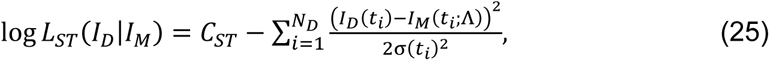

where *σ(*t*)* is approximated by the measured SEM and *N_D_* is the number of time points measured in the drug-treatment curve, and *C_ST_* is constant that doesn’t depend on model parameters.

For chemical stress experiments, three data sets were used representing the intensity for Cap-only spots, IRES-only spots, and green (Cap) intensity in both Cap and IRES spots. These three data sets were considered on a total log-likelihood function as follows:

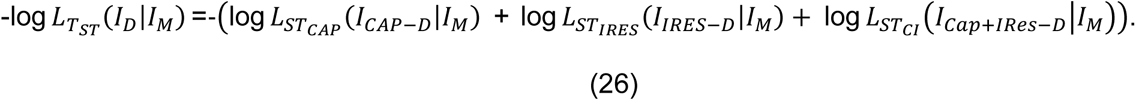

### Uncertainty Quantification

To quantify uncertainty, the best parameter set from fitting was initially used and then a Markov Chain Monte Carlo (MCMC) algorithm was run to explore an additional 10,000 possible parameter combinations. At each step, a random perturbation to the current parameters was proposed and every proposal for which the log-likelihood for the new parameter set was within a 1% of that found for the best fit was accepted (i.e., all parameters for which log*(L(*I*_D_|*I*_Best_)/L(*I*_D_|*I*_New_)) < 60* were accepted). The resulting parameter sets were then used to estimate standard deviations of the parameters as shown in Table 1.

### Computational Implementation and Codes

All simulations were performed on the W. M. Keck High Performance Computing Cluster at Colorado State University. All codes and required data will be available in the following repository: https://github.com/MunskyGroup/CAP_IRES.

---

**Figure S1:**
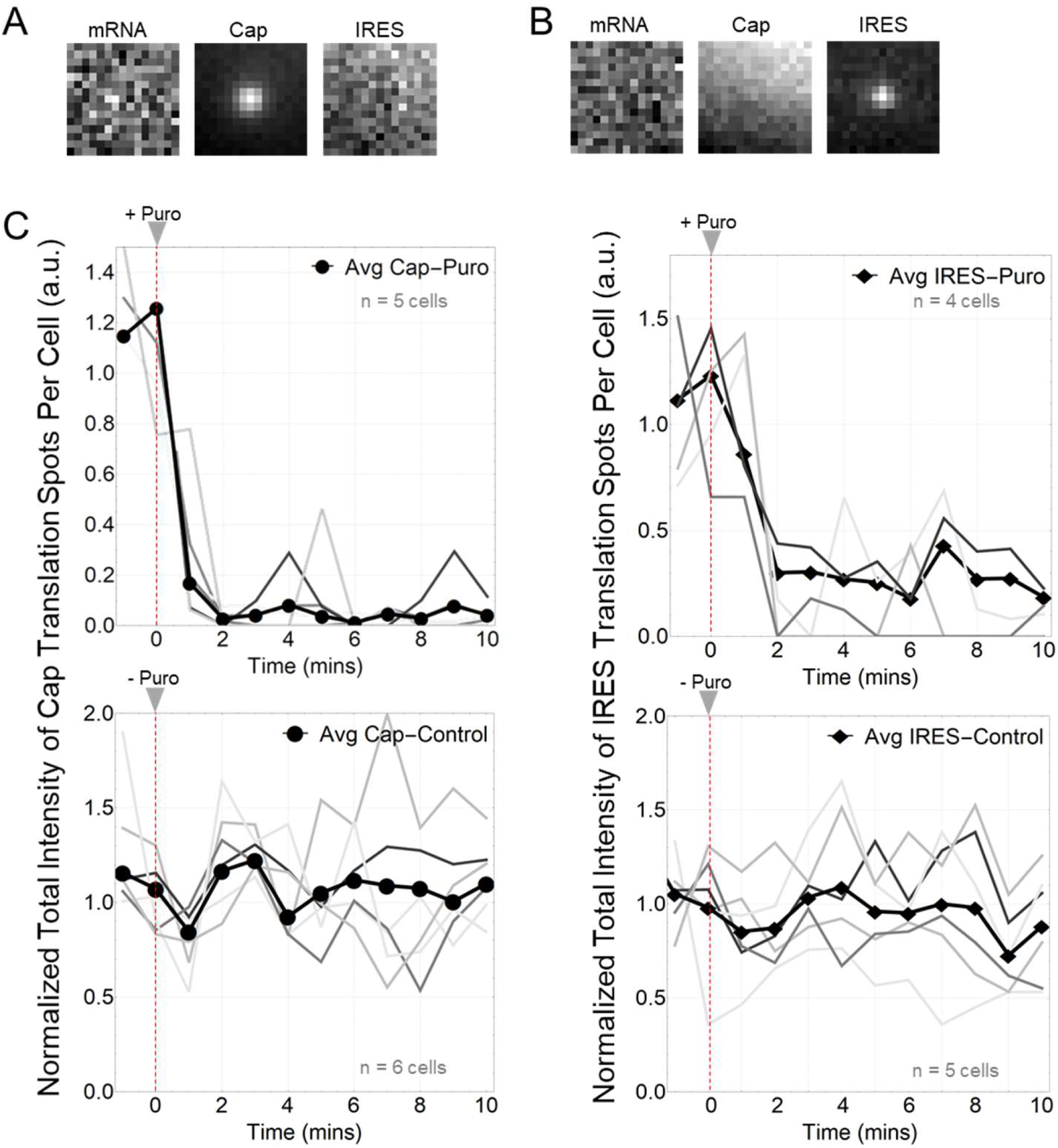
Controls for bleedthrough and active translation. (A-B) Five frame average of a Cap Only and IRES Only translation spots. mRNA marker dye, JF646, was not added to these cells. Cells were imaged for 3-minutes with a 6-second interval between each capture. (C) Top graphs show normalized total intensity over time for Cap-dependent (left) and IRES-mediated translation spots (right), after addition of puromycin. Gray lines indicate individual cells. Thick dark line indicates the average total intensity of all cells. Red dashed lines indicate time at which puromycin was added. Cap-dependent: n=5 cells. IRES-mediated: n= 4 cells. Bottom graphs show normalized total intensity of Cap-dependent (left) and IRES-mediated (right) translation spots without the addition of puromycin. Cap-dependent: n=6 cells. IRES-mediated: n= 5 cells. All cells (control and drug treated) were imaged for 10-minutes with a 1-minute interval.

**Figure S2:**
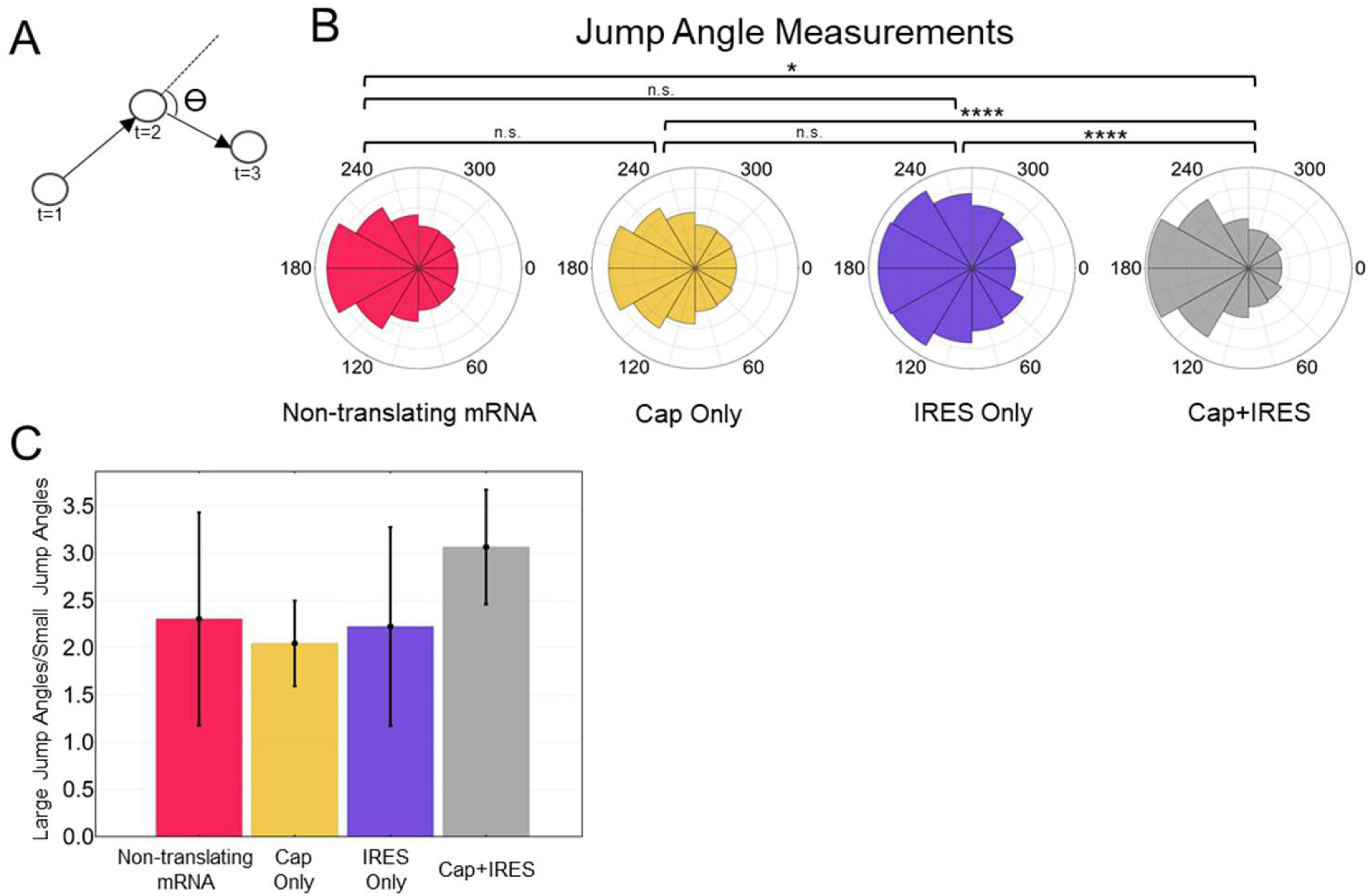
Jump angle measurements to quantify confinement in non-translating and translating mRNA. (A) Schematic showing how the jump angles are measured. (B) Circular plots of the jump angle distributions for non-translating mRNA, Cap Only, IRES Only and Cap+IRES translation sites. (C) Bar plots showing the ratios of large jump angles (150-180 degrees) to small jump angles (0-30 degrees) for non-translating mRNA, Cap Only, IRES Only, and Cap+IRES translation sites. Angles are measured in degrees. The p-values are based on a two-tailed Mann-Whitney test: *p<0.05, **p<0.01, ***p<0.001, ****p<0.0001.

**Figure S3:**
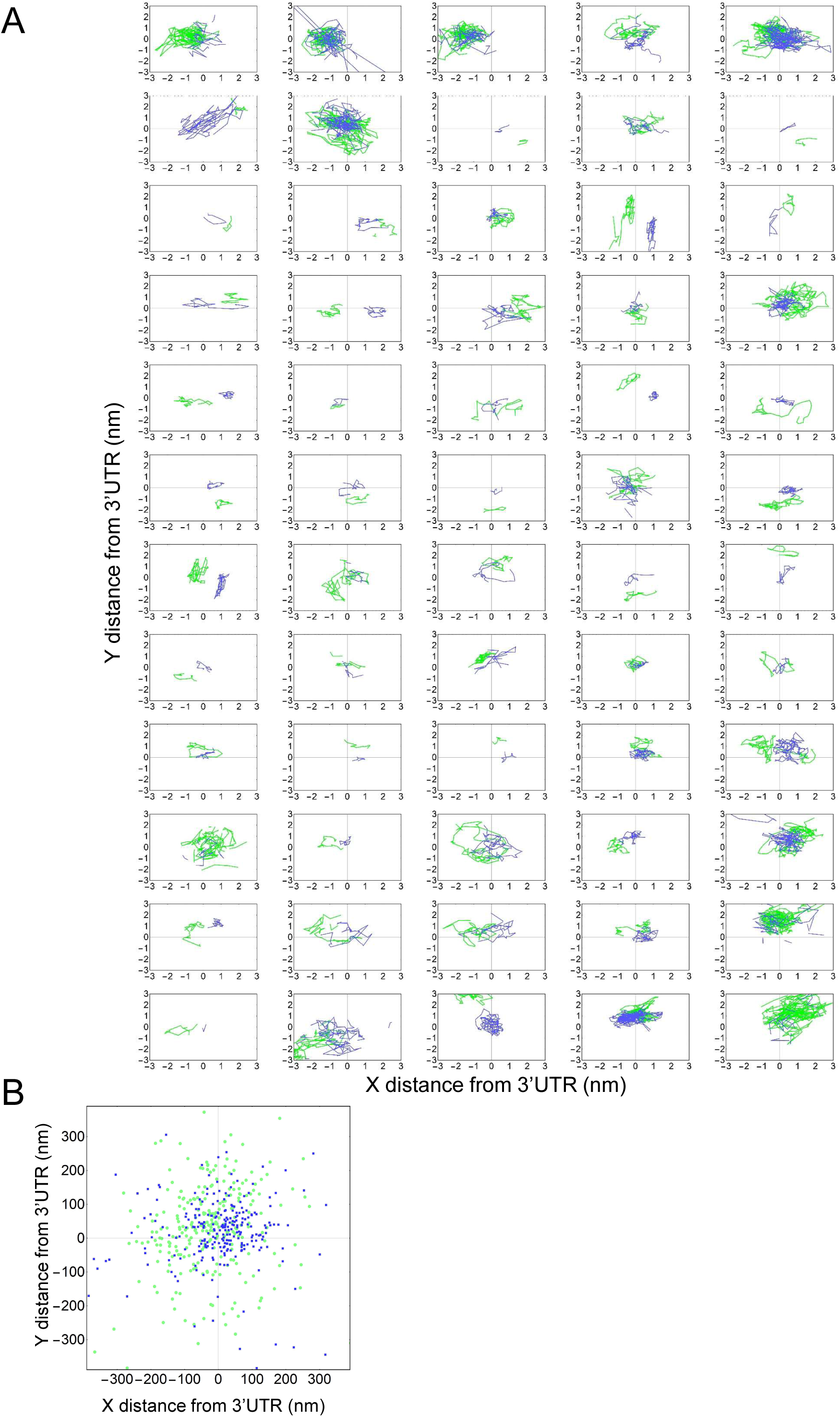
Measuring distances between Cap and IRES nascent chains in Cap+IRES translation spots. (A) Representative data set of measured distances of Cap (light green) and IRES (blue) nascent chains to 3’UTR through time in single Cap+IRES translation tracks. (B) Median distances of Cap and IRES nascent chains to 3’UTR of each Cap+IRES track. Distances are measured in nanometers (nm). 3’ UTR coordinates were fixed at (0,0) for all analyses.

**Figure S4:**
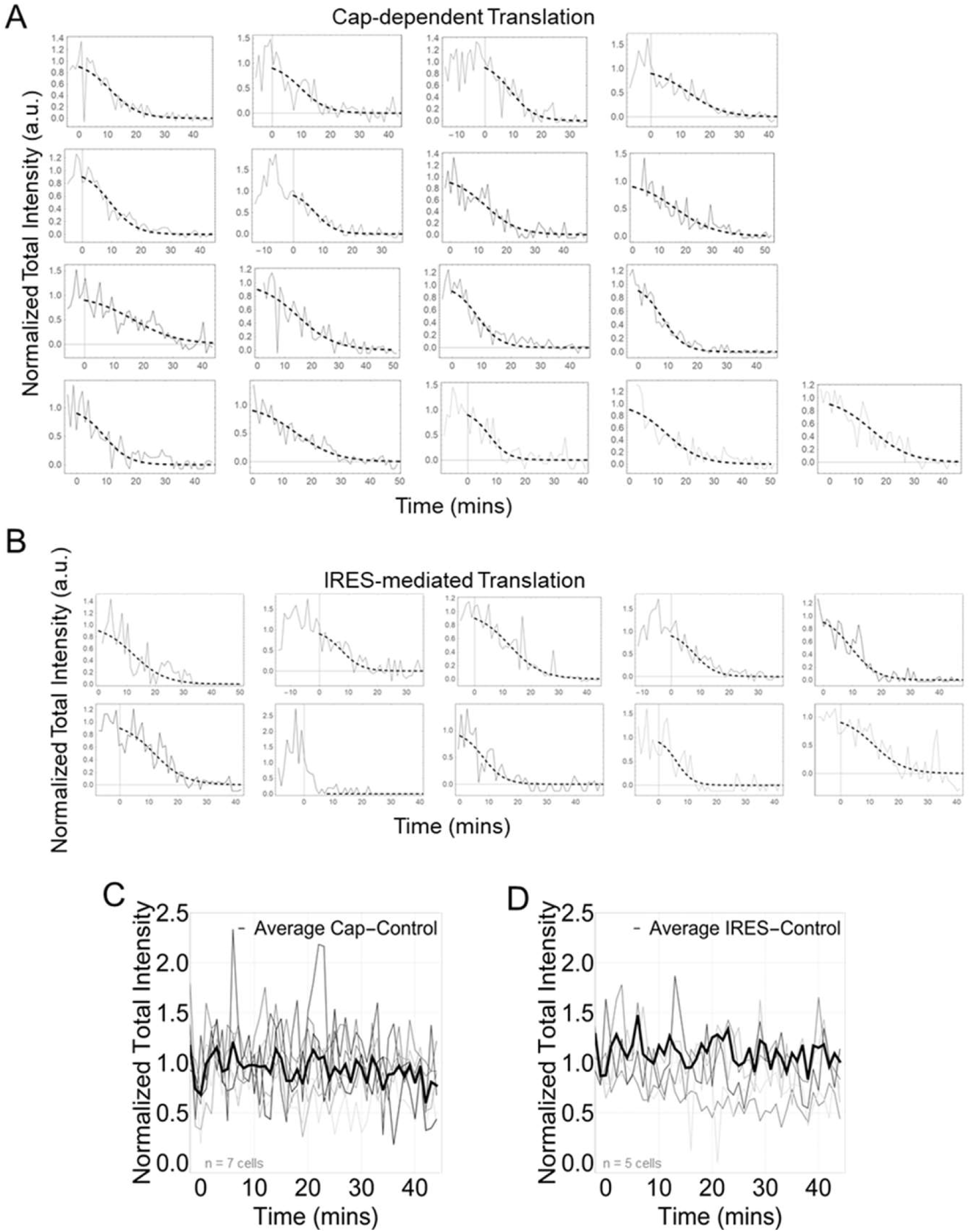
Ribosomal run-off curves from single cells after addition of Harringtonine. (A) Harringtonine-induced ribosomal run-off curves from single cells. Each curve shows the decay in nascent chain signal intensity from all Cap-dependent and (B) IRES-mediated translation sites within a single cell post-Harringtonine. Run-off curves were phenomenologically fit to a Tanh function to align curves in time for averaging in Figure 4. The slope of fitted curves at a normalized intensity value of 0.5 was used to estimate the elongation rate. (C) Cap-dependent (n=7 cells) and (D) IRES-mediated (n=5 cells) translation controls in which no drugs were added. Each gray line shows the total nascent chain signal intensity from all translation sites in an individual cell. The thick black line represents the average intensity from all cells. Intensity in arbitrary units (a.u.). All cells were imaged for 45 minutes with a 1-minute interval between each capture.

**Figure S5:**
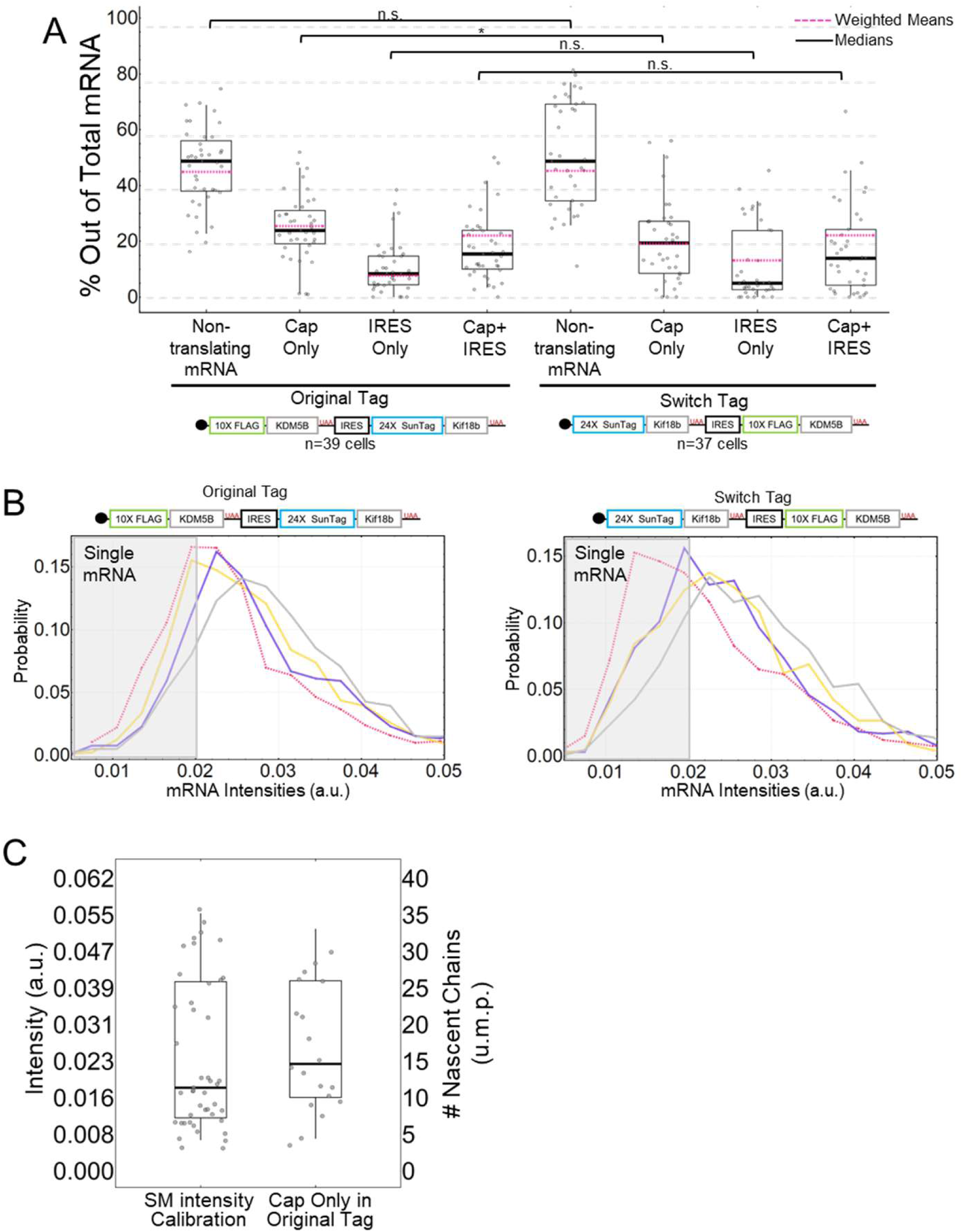
Original Tag comparison to Switch Tag, single mRNA selection, and polysome intensity calibrations. (A) Quantification of the percentages of each type of translation sites for the Original Tag (left, n=39 cells) and the Switch Tag (right, n=37 cells). Each point represents a single-cell measurement. The p-values are based on a two-tailed Mann-Whitney test: *p<0.05, **p<0.01, ***p<0.001, ****p<0.0001. The thick black lines indicate the median, the dashed red line represents the weighted mean, the boxes indicate the 25%-75% range, and the whiskers indicate the 5%-95% range. (B) Probability histograms showing distributions of mRNA intensities of non-translating mRNA (Red), Cap Only (Yellow), IRES Only (Purple), and Cap+IRES (Gray) translation sites for the Original Tag and the Switch Tag. The gray boxes represent the mRNA intensity threshold used to eliminate multiple mRNAs. Intensities in arbitrary units (a.u.). (C) Translation site calibration measurements. The intensities of Cap in Cap Only translation sites (n=20spots) in the Original Tag were compared to a 10xFlag calibration system (n=47spots) with a known number of ribosomes. These comparisons lead to a calculated number of 14.6 ribosomes in Cap Only translation sites using the Original Tag.

**Figure S6:**
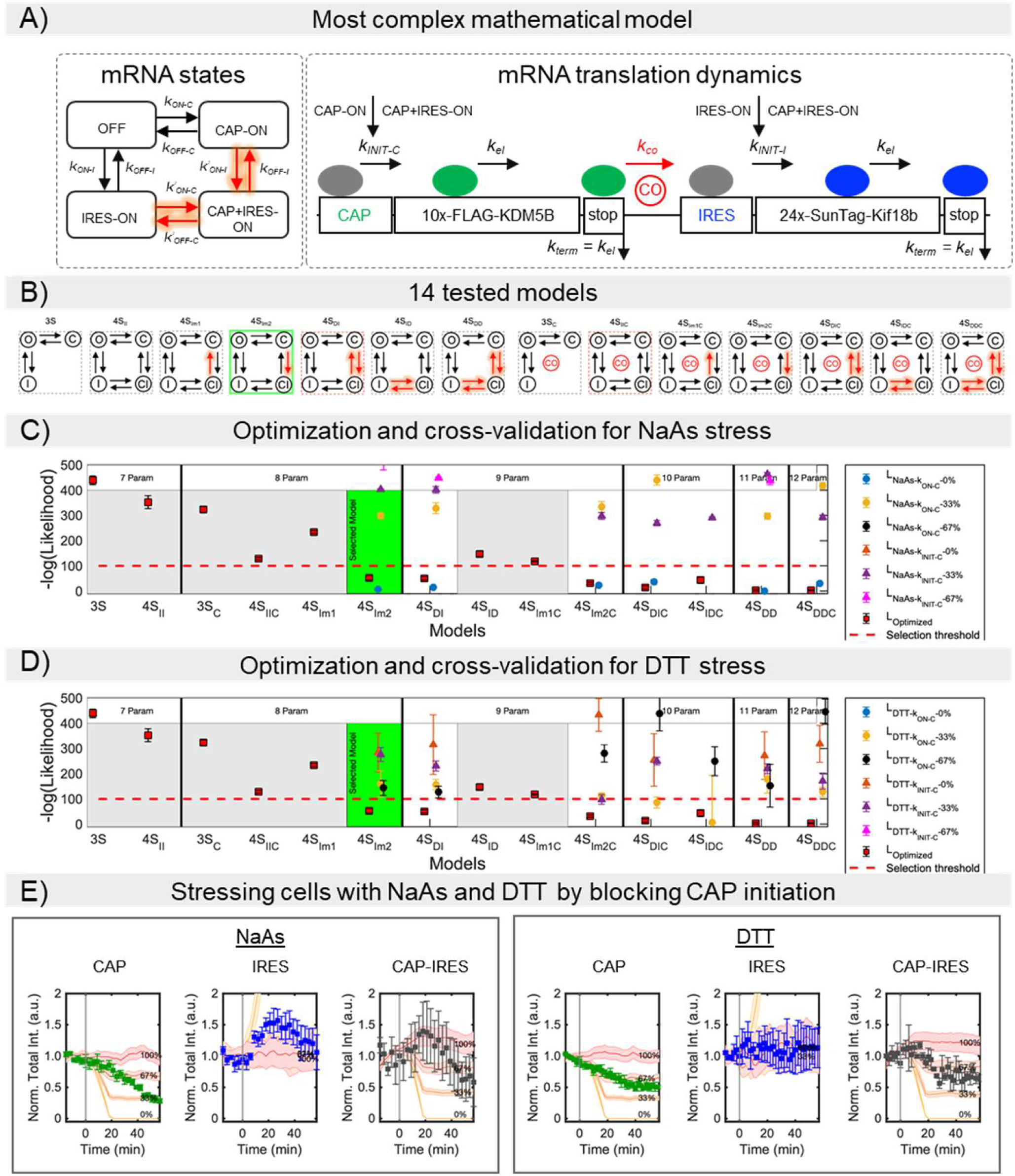
Model of the bicistronic gene construct. (A) The most complete mathematical model considers four mutually exclusive RNA states: non-translating (OFF), Cap-dependent (CAP-ON), IRES-mediated (IRES-ON), and both Cap and IRES (CAP+IRES-ON). All transition rate values between RNA states are free-independent parameters. A cross-over mechanism (CO symbol in the figure), by which a ribosome that completes the translation of the Cap-dependent protein could immediately re-initiate translation of the IRES-mediated protein, is represented by the reaction parameter *k_co_*. (B) Comparison of 14 different sub-models. The sub-models test different hypotheses, including variations of the number of mRNA states (3 or 4 states), dependency on Cap and IRES switching states, and/or the existence of the cross-over mechanism. Cap and IRES dependency are represented in the figure by red lines, which denote that the corresponding reaction parameter value has a free value during the optimization process. All models have 3 or 4 mRNA states, denoted by 3S or 4S, respectively. From left to right, the first seven models lack crossover, while the last seven have cross-over (denoted by subscript *‘C*’, e.g. 3S_C_). Models can have independent (denoted by subscript ‘*I*’) or dependent (denoted by subscript ‘*D*’) Cap or IRES activation/deactivation. Models can also have a single dependent activation or deactivation rate (denoted by subscript ‘*m1*’ or ‘*m2*’). The number of free parameters in the sub-models ranges from 7 to 12. (C) Cross-validation is used to compare two possible mechanisms of translation inhibition under DTT stress. The first mechanism mimics the inhibition of the Cap activation rates at the promoter level (*L_NaAs-STATE-CAP_*; i.e., block of *k_ON-C_* and *k’_ON-C_*). The second mechanisms considers blocking ribosomal initiation for Cap (*L_NaAs-INIT-CAP_*; i.e., block of *k_INIT-C_*). (D) Optimization process and cross-validation for the DTT stress. The same inhibitory mechanisms described in C are tested for DTT stress. Relative Log-likelihood values for the optimization process are calculated according to Eq. 23 and according to Eq. 26 for the NaAs and DTT cross-validation experiments, respectively. The log-likelihood reported are relative to the minimum value from all models. Relative log-likelihood values over 500 are not plotted. A selection threshold (dashed red line) was defined by a log-likelihood of 100 worse than the most complex and best fitting model. Models above the selection threshold were discarded (gray background), and their cross-validation log-likelihood values are not shown. The best model shown (green background) was chosen as the model with fewest free parameters below the selection threshold. (E) Model simulations for the best-fit model *4S_Im2_* under under NaAs and DTT stresses. The figure shows the effect of blocking ribosomal initiation for Cap. The assumption of blocking initiation results in significantly worse predictions compared to the hypothesis that the drugs block Cap activation (compare to Fig. 6F).

